# Rapid evolution of gene expression patterns in flowering plants

**DOI:** 10.1101/2024.07.08.602577

**Authors:** Christoph Schuster, Alexander Gabel, Hajk-Georg Drost, Ivo Grosse, Ottoline Leyser, Elliot M. Meyerowitz

**Author notes:** These authors contributed equally to this work (Alexander Gabel, Hajk-Georg Drost). Correspondence: Christoph Schuster, Elliot M. Meyerowitz, (C.S.), (E.M.M.).

## Abstract

Phenotypic differences between species are largely driven by changes in both protein-coding sequence and gene expression ^1^. The evolutionary history of angiosperms (flowering plants) is characterised by a highly accelerated rate of diversification, which Darwin referred to as an “abominable mystery” ^2^. Here we show, by analysing the transcriptomes from eight organs across seven species, that angiosperm protein-coding gene expression patterns evolve rapidly: within 45 million years, expression levels of orthologous genes diverged so strongly that they are more similar between different organs within a species than between what are considered homologous organs from different species. This finding differs from previous observations in mammals, which demonstrated that organ-dependent gene expression levels are largely conserved ^3, 4, 5^. Among the angiosperm organs, meristems and leaves show the highest degree of expression conservation, whereas stamen and pollen transcriptomes diverge rapidly. Examining changes in the expression level of functionally related genes, we found low rates for those involved in key cellular, metabolic and developmental processes. In contrast, particularly high rates were observed for genes that are involved in the response to endogenous and external stimuli, presumably reflecting an adaptive response of flowering plants to ever-changing environments. Our work reveals that the evolution of gene expression progresses at different rates in angiosperms and mammals, and provides a comprehensive resource to perform cross-kingdom comparative studies of transcriptome evolution.

---

Whole-genome hominid evolutionary studies have shown that phylogenetically closely related species, despite exhibiting substantially distinct morphological phenotypes, share a large proportion of protein-coding genes, suggesting that alterations in coding sequence only partially explain phenotypic innovation ^6, 7^. Indeed, changes affecting the spatio-temporal expression of genes largely contribute to the phenotypic evolution of species and have been extensively studied in mammals in recent years ^3, 4, 5, 8^. Given the relatively low rate of diversification of mammalian species compared to the overwhelming number and ecological dominance of angiosperms (6649 mammalian *versus* > 300,000 angiosperm species) ^9, 10^, we hypothesised that gene expression levels may have evolved at different rates in both clades and kingdoms. To test this prediction, we generated a comprehensive total RNA-seq bulk transcriptome dataset of eight organs from seven angiosperm species. By combining our dataset with available data from mammals ^3^, we explore how the evolution of both coding and noncoding gene expression in flowering plants compares to the evolutionary rates found in animals.

## Angiosperm organ transcriptomes

High-throughput whole transcriptome sequencing (RNA-seq) was conducted using triplicate ribo-depleted total RNA samples from root, hypocotyl, leaf, vegetative apex, inflorescence apex, flower, carpel, and stamen, using *Arabidopsis thaliana* (Col-0) as the reference species. We also collected RNA-seq data from mature pollen as a representative tissue consisting of only two cell types, a generative cell and one or two sperm cells, allowing us to analyse changes in gene expression between species at almost cell-type-specific resolution. Species were chosen based on the quality of their published genome sequence and their evolutionary distance, consisting of the four closely related Brassicaceae species *Arabidopsis thaliana*, *Arabidopsis lyrata*, *Capsella rubella*, and *Eutrema salsugineum*; *Tarenaya hassleriana* from the Brassicaceae sister family Cleomaceae, the legume *Medicago truncatula*, and the monocot *Brachypodium distachyon* (Fig. 1a). In addition to these comparative samples, we prepared transcriptome libraries from a wide range of tissues and developmental stages of *A. thaliana*, from early embryogenesis to senescence, to study patterns of coding and non-coding gene expression at higher spatio-temporal resolution. The complete dataset, which we termed DevSeq gene expression atlas, consists of 303 libraries (Extended Data Table 1, Extended Data Fig. 1a-c, i). Based on protein sequence similarity, we identified a set of 7003 protein-coding 1–1 orthologous genes for comparisons across all species. Most of the long non-coding RNAs (lncRNAs) we detected were species-specific, with only 307 lncRNA orthologs shared between the four Brassicaceae species, and only 8 identifiable across all seven species (Fig. 1b).

**Figure 1.**
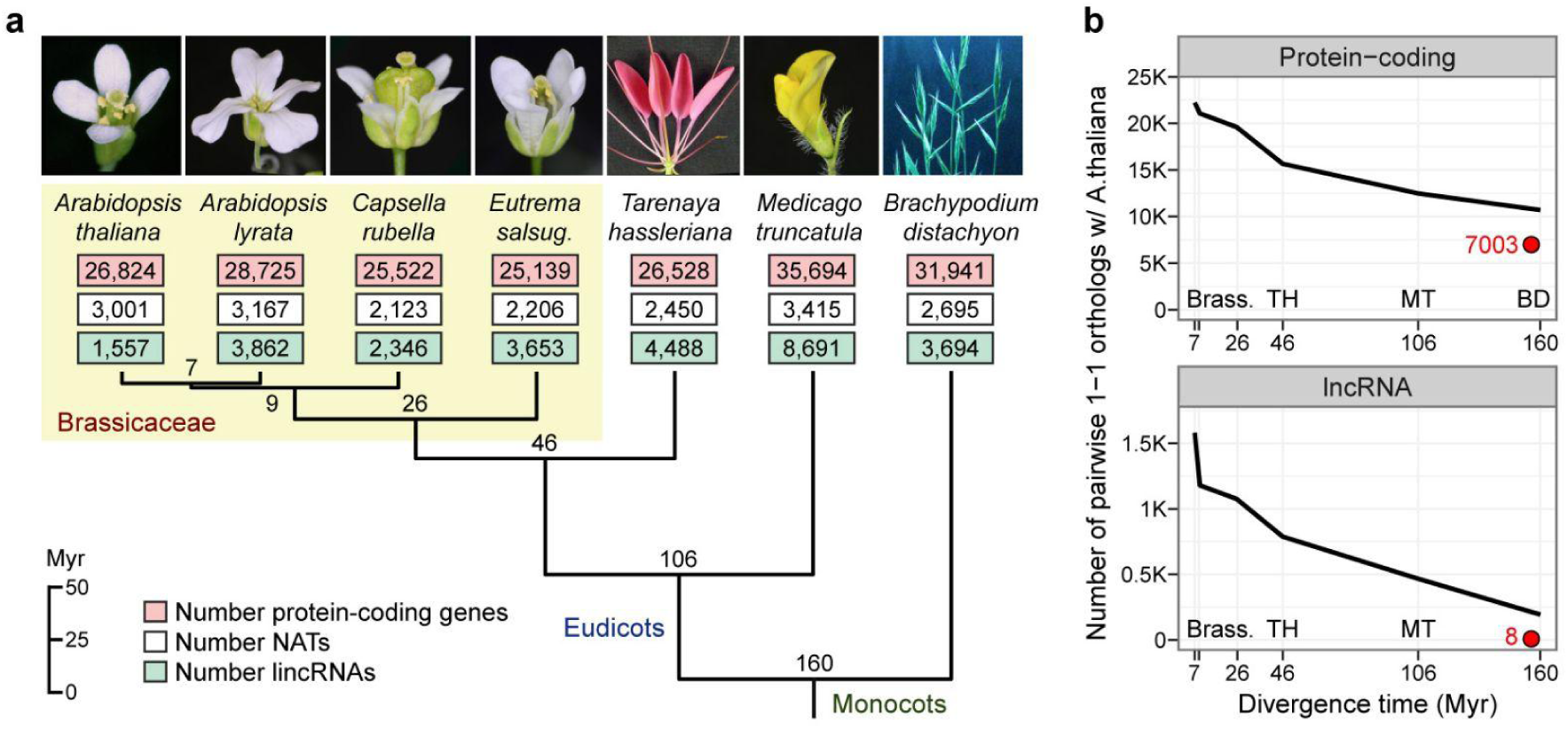
Conservation of protein-coding genes and lncRNAs in angiosperms. **a,** Phylogenetic tree of angiosperm species used in this study. Numbers on nodes indicate TimeTree of Life (TTOL) ^11^ estimated divergence times between species in million years (Myr). Tree tips show numbers of protein-coding genes, cis-natural antisense transcripts (NATs) and long-intergenic non-coding RNAs (lincRNAs) for each species. **b,** Number of pairwise protein-coding and long non-coding RNA (lncRNA) 1-1 orthologs with *Arabidopsis thaliana*. Red circles indicate the number of orthologous protein-coding gene and lncRNA families identifiably conserved across all species. Brass., Brassicaceae; TH, *Tarenaya hassleriana*; MT, *Medicago truncatula*; BD, *Brachypodium distachyon*. Divergence time, TTOL estimated times.

## Gene expression patterns across organs and species

Depending on the respective plant species, we identified approximately 25,000 to 35,000 protein-coding genes, 2,100 to 3,400 cis-natural antisense transcripts (cis-NATs), and 1,500 to 8,600 long-intergenic non-coding RNAs (lincRNAs) being transcribed above the sample-specific, minimal spike-in threshold (Fig. 1a). Overall, different plant organs show a similar number of expressed protein-coding genes and non-coding RNAs in all species, and substantially fewer transcripts were detected in pollen, reflecting the unique transcriptome characteristics of the male gametophyte and the low number of cell types (Extended Data Fig. 1d-g and Extended Data Fig. 2a-d) ^12, 13, 14^. Consistantly, hierarchical clustering of transcriptome profiles from *Arabidopsis thaliana* reveals pollen and stamen as the primary outgroup, and recapitulates organ type, highlighting the distinct expression signatures of individual organs (Extended Data Fig. 1h).

A more detailed analysis of samples from the reference plant *Arabidopsis thaliana* covering all major organs and many developmental stages showed that most protein-coding genes exhibit maximum expression intensity in root tips, stamens at stage 12 and mature seeds, and lncRNAs are preferentially expressed in stamens at stage 12 (Fig. 2a,b and Extended Data Fig. 2e,f). These patterns are consistent across different conservation levels of coding and non-coding genes, and are found in all other angiosperm species used in this study, except *Tarenaya hassleriana* (Extended Data Fig. 3a,b). In addition, we noticed that highly conserved protein-coding genes (core orthologs; minimum age 160 Myr) show higher maximum expression intensities than young genes (Fig. 2c,d). This observation is consistent with previous studies in vertebrates and yeast, which demonstrated that the intensity of gene expression inversely correlates with the rate of protein sequence evolution ^15, 16^. A similar trend was observed when comparing the maximum expression of non-conserved and Brassicaceae-conserved (minimum age 26 Myr) lncRNAs (Fig. 2c,d).

**Figure 2.**
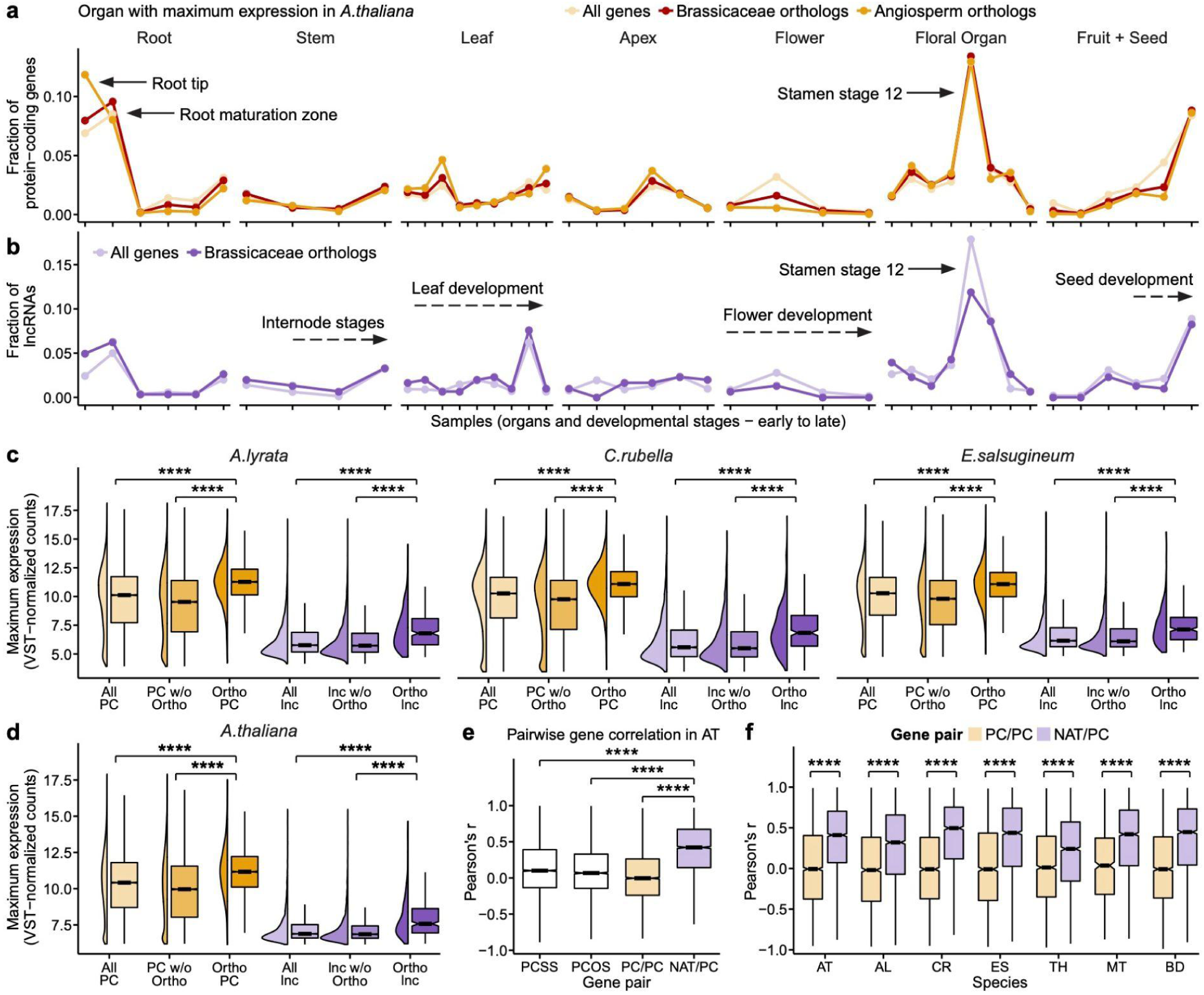
Expression patterns of coding and non-coding genes in angiosperms. **a-b,** Fraction of protein-coding genes (**a**) and long non-coding RNAs (lncRNAs) (**b**) for which maximum expression was detected across 43 *Arabidopsis thaliana* sample types. Sample types (x-axis) were assayed in pooled biological triplicates and represent different zones and developmental stages of all major organs. Lines indicate the proportion of all expressed protein-coding genes (n = 26,824), Brassicaceae protein-coding orthologs (n = 17,445), and angiosperm protein-coding (core) orthologs (n = 7,003) (**a**), and the proportion of all lncRNA genes (n = 4,693) and Brassicaceae lncRNA orthologs (n = 307) (**b**). **c-d,** Distribution of the maximum expression level of all detected protein-coding genes, non-orthologous protein-coding genes, angiosperm protein-coding orthologs, all lncRNAs, non-orthologous lncRNAs, and Brassicaceae orthologous lncRNAs across eight organs (root, hypocotyl, leaf, apex vegetative, apex inflorescence, flower, stamen, carpel) in the four Brassicaceae species studied. **e-f,** Co-expression of protein-coding genes and cis-natural antisense transcripts (cisNATs) in *Arabidopsis thaliana* (**e**), and in the other species (**f**). Shown are neighbouring, non-overlapping protein-coding genes located on the same strand (PCSS) or on opposite strands (PCOS), overlapping protein-coding gene pairs (PC/PC) and cisNAT/protein-coding gene pairs (NAT/PC). **** *P* < 10-6, two-sided Wilcoxon rank-sum test.

While earlier studies in *Arabidopsis thaliana* based on massively parallel signature sequencing ^17^ and microarray data ^18^ provided evidence for an antagonistic activity pattern between cis-NATs and their cognate sense protein-coding genes (PC), we found that the expression profiles of the majority of cis-NAT/PC pairs are positively correlated. In contrast, the expression of neighbouring, protein-coding genes that do not overlap was only very weakly correlated ^19^, and no relationship was observed for overlapping protein-coding gene pairs (Fig. 2e). This trend was found across all species (Fig. 2f). Further analysis provided no evidence for a relationship between cis-NAT expression intensity, cis-NAT/PC expression ratio nor cis-NAT/PC overlap length and the expression correlation of the cis-NAT/PC pair (Extended Data Fig. 3c-e). Recently, several mechanisms that contribute to the crosstalk between intergenic lncRNAs and neighbouring genes have been described that do not require the lncRNA transcripts themselves, but instead depend on the processes related to their production. These include the process of transcription as such by recruiting activating factors or remodelling nucleosomes, and enhancer-like activity of gene promoters ^20, 21^. It is tempting to speculate that cisNATs might influence the expression of their cognate sense transcripts in similar ways.

To further explore to what extent technical aspects of expression quantification, sample handling, differences in growth conditions and variations in tissue dissections influence the overall results of downstream analyses, we compared our data set with one of the most comprehensive *Arabidopsis thaliana* developmental transcriptome maps to date, the AtGenExpress gene expression atlas ^13^. In total, 26 samples common to both data sets, collected from organs at the same or a comparable developmental stage, were included in this analysis. Hierarchical clustering of pairwise Pearson distances revealed that the samples clustered by organ rather than by study (Extended Data Fig. 4a), and dendrograms of samples from both studies looked remarkably similar (Extended Data Fig. 4c-d). In addition, individual gene expression profiles across organs and developmental stages were highly correlated between studies (Extended Data Fig. 4b,e). These results illustrate that our experimental setup is both accurate and robust enough to detect the unique expression signatures of samples across development, despite differences in growth conditions, dissections, library construction protocols and quantification platforms.

## Rapid evolution of gene expression levels in angiosperm organs

The main goal of the project was to analyse the evolutionary dynamics of coding and non-coding gene expression in angiosperms. To investigate the relationship between samples, we first performed hierarchical clustering based on Pearson distances calculated from the expression values of 7003 orthologous protein-coding genes present in all seven species. While samples of the same organ from closely related species partially clustered together, we found that samples from distant angiosperms (*T. hassleriana*, *M. truncatula*, *B. distachyon*) group invariably by individual species (Fig. 3a and Extended Data Fig. 5b). This result differs from the findings of recent studies in mammals, which demonstrated that even organs from distantly related species, such as primate species and mouse, group together, suggesting that the conservation of core regulatory networks and developmental pathways in organs limit transcriptome divergence ^3, 22^. Since it has been shown that this tissue-dominated clustering pattern largely depends on the organs being compared ^23^, we asked whether the angiosperm organs we chose were more similar to each other than those evaluated in previous mammalian studies. Thus, to be directly comparable we re-analysed the data from Brawand *et al.* ^3^ using our ortholog detection and RNA-seq data analysis pipeline. As previously reported, mammalian samples from the same tissue from different species cluster together, with the exception of brain and cerebellum, which have been reported to possess similar expression signatures ^3, 4^ (Extended Data Fig. 5a). We calculated intraspecies Pearson distances among tissues and consistently confirmed brain-cerebellum as the sample pair with the smallest distance. When plotting the distribution of intra-species distances among angiosperm organs, we found that they are on average more similar compared to organs from the mammalian dataset. We identified vegetative apex-inflorescence apex, flower-stamen and flower-carpel as the organ pairs with the smallest distance (Extended Data Fig. 5d). After excluding cerebellum samples from the mammalian data set, and apex tissues and flower samples from the angiosperm data set, the resulting mammalian and angiosperm subsets revealed a very similar distribution of expression distances (Extended Data Fig. 5e). However, angiosperm organs from the phylogenetically distant species still grouped by species rather than organ, indicating that gene expression patterns of orthologous genes from different organs of the same species are more similar than signatures of what are considered homologous organs from distant species. This species-dominated clustering pattern of distant angiosperms was further investigated by principal component analysis (PCA). Reducing the expression data of 17,445 orthologous protein-coding genes present in all four Brassicaceae genomes to its essential features, we observed that data separated by organ (Fig. 3b, Extended Data Fig. 5f-h). In contrast, this pattern was less evident when mapping the major components of the expression data from 7,003 angiosperm orthologs using the complete data set. Here, the distances between species were of the same order of magnitude as those between organs, and the phylogenetically most distant species were separated from the rest (Fig. 3c, Extended Data Fig. 5i-k). In order to better understand the relative rates at which protein-coding gene expression has diverged, we calculated pairwise Pearson correlations between what are considered homologous organs from *Arabidopsis thaliana* and the other species, based on the expression values of the angiosperm orthologs (Fig. 3d). Whereas the organ transcriptomes remained rather similar within the Brassicaceae family, they rapidly diverged between *Arabidopsis thaliana* and the evolutionarily more distant species, in line with the results of the clustering analysis. We also calculated Pearson correlations using *Arabidopsis lyrata* as reference species and obtained a comparable result (Extended Data Fig. 6h).

**Figure 3.**
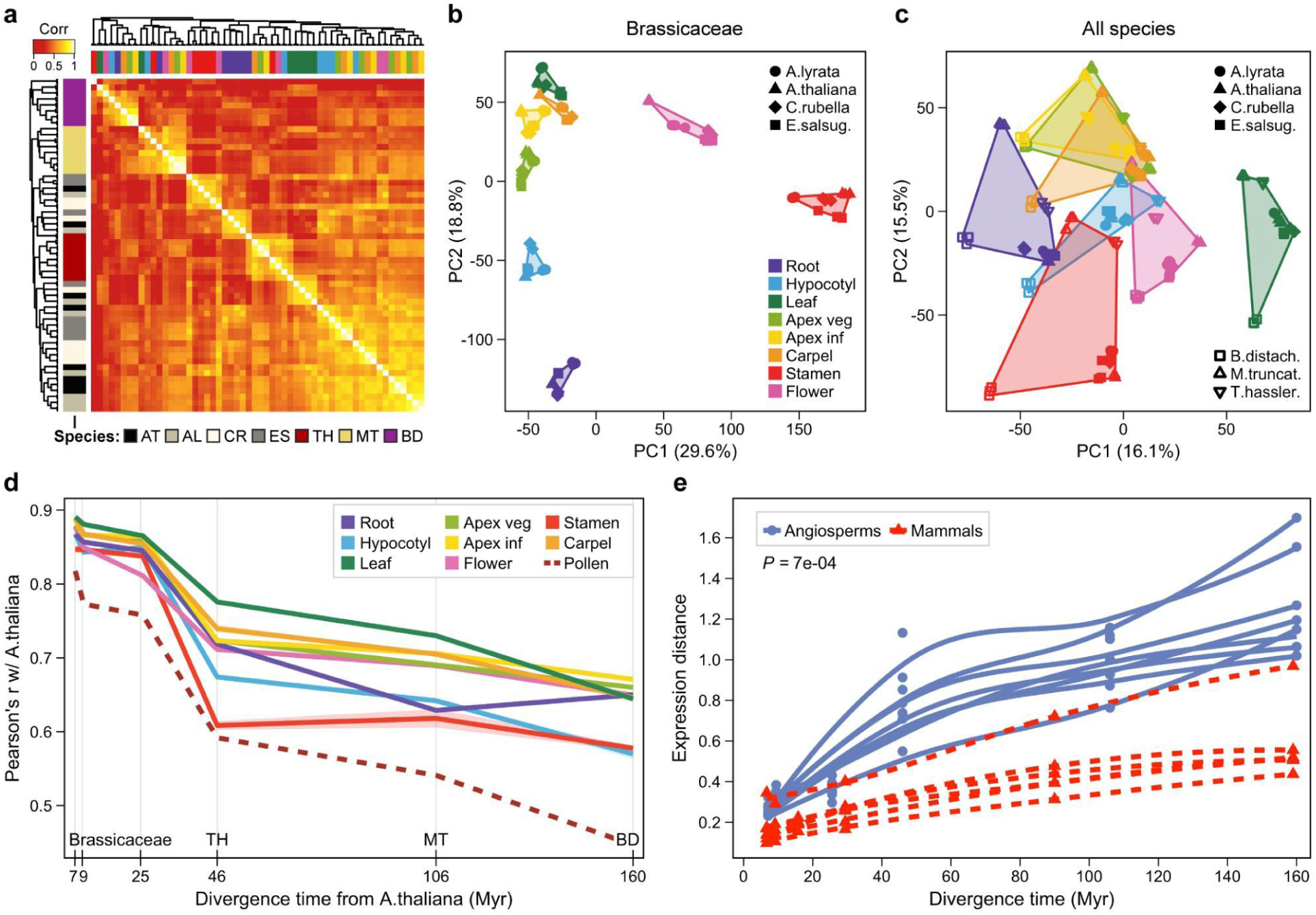
Gene expression levels evolve rapidly in angiosperms. **a,** Symmetrical heat map of Pearson correlations from angiosperm organ transcriptomes. Expression levels of 7,003 1-1 orthologous protein-coding genes, estimated as normalized counts after variance-stabilizing transformation (VST), were analysed across 8 organs and 7 species. Samples were hierarchically clustered based on Pearson distance using complete linkage method. Colour bars: horizontal, organs; vertical, species. Organ legend is shown in (b). AT, *Arabidopsis thaliana*; AL, *Arabidopsis lyrata*; CR, *Capsella rubella*; ES, *Eutrema salsugineum*; TH, *Tarenaya hassleriana*; MT, *Medicago truncatula*; BD, *Brachypodium distachyon*. **b, c,** Principal-component analysis of protein-coding gene expression levels from 17,445 1-1 orthologs in Brassicaceae (**b**), and from 7,003 1-1 orthologs conserved across all analysed angiosperm species (**c**). **d,** Pairwise Pearson correlations between *Arabidopsis thaliana* and the other species, for 8 organs and pollen, based on mRNA gene expression levels (log-transformed TPM, 7,003 genes). For *Brachypodium*, mesocotyl was analysed instead of hypocotyl. **e,** Expression distance estimated under the Ornstein-Uhlenbeck (OU) model, in angiosperms and mammals {Brawand, 2011 #402}. Each point corresponds to the average distance between all species pairs across the entire phylogenetic tree, for each individual organ. For angiosperms, expression distances were calculated for 8 organs from 7 species (same as in a, c, d, without pollen). Divergence-time estimates are 7, 9, 26, 46, 106, and 160 Myr. Mammalian expression distances were calculated for 6 organs (brain, cerebellum, heart, kidney, liver, and testis) from 8 species: *Homo sapiens*, *Pan troglodytes* (chimpanzee), *Pan paniscus* (bonobo), *Gorilla gorilla, Pongo pygmaeus* (orangutan), *Macaca mulatta* (macaque), *Mus musculus* (mouse), and *Monodelphis domestica* (opossum); divergence time estimates are 7, 9, 16, 29, 90, and 159 Myr. Lines show expression divergence of individual organs and were estimated by LOESS regression. Organ gene expression levels diverge faster in angiosperms than in mammals (Ρ = 7 x 10^-4^, two-sided Wilcoxon rank-sum test).

While the above findings illustrate that protein-coding gene expression levels are conserved among closely related angiosperm species and have evolved rapidly over longer evolutionary distances, they did not allow direct quantification of the rate of transcriptome evolution among angiosperm and mammalian organs. To this end, we retrieved two measures of expression distances for each organ: the Pearson distance, preserved as a metric, and an expression distance estimated under the Ornstein-Uhlenbeck model ^24^. Both distances were calculated between all species pairs across the entire phylogenetic tree. We then applied two different non-linear regression methods, a negative exponential growth model and locally estimated scatterplot smoothing (LOESS), and determined the average slope value as a measure of transcriptome evolution (Extended Data Fig. 6a-f). Consistent with earlier studies in *Drosophila* ^25^ and mammals ^26^, we found that transcriptome divergence saturates over evolutionary time, indicating that gene expression levels are dominantly under stabilising selection (Extended Data Fig. 6a-c). To evaluate possible differences in the rate of expression evolution between angiosperm and mammalian organs, we compared the average slope values from the non-linear regression fits. For both distance metrics, angiosperm organs exhibit higher slope values than mammalian organs, confirming our initial hypothesis that gene expression levels evolved at a different rate in both clades (Ρ = 7 x 10^-4^, two-sided Wilcoxon rank-sum test, Fig. 3e, Extended Data Fig. 6g). To further analyse the evolutionary changes in gene expression for each individual organ, we constructed neighbour-joining trees based on pairwise Pearson distances between species. The reliability of the branches in the phylogenetic trees was assessed by randomly re-sampling the 1–1 orthologous genes with replacement (7,003 1–1 orthologs, 1,000 bootstraps). Notably, the gene expression phylogenies largely followed the taxonomic relationships, with some exceptions only between closely related members of the Brassicaceae family. The distribution of the total branch lengths from the bootstrap trees display differences between organs; meristematic organs (vegetative and inflorescence apex, carpel) and leaf have the smallest branch lengths, indicating low rates of expression evolution. In contrast, stamen and pollen diverged most rapidly (Fig. 4a). Similar results were observed when we limited the analysis to the Brassicaceae data set (Fig. 4b).

**Figure 4.**
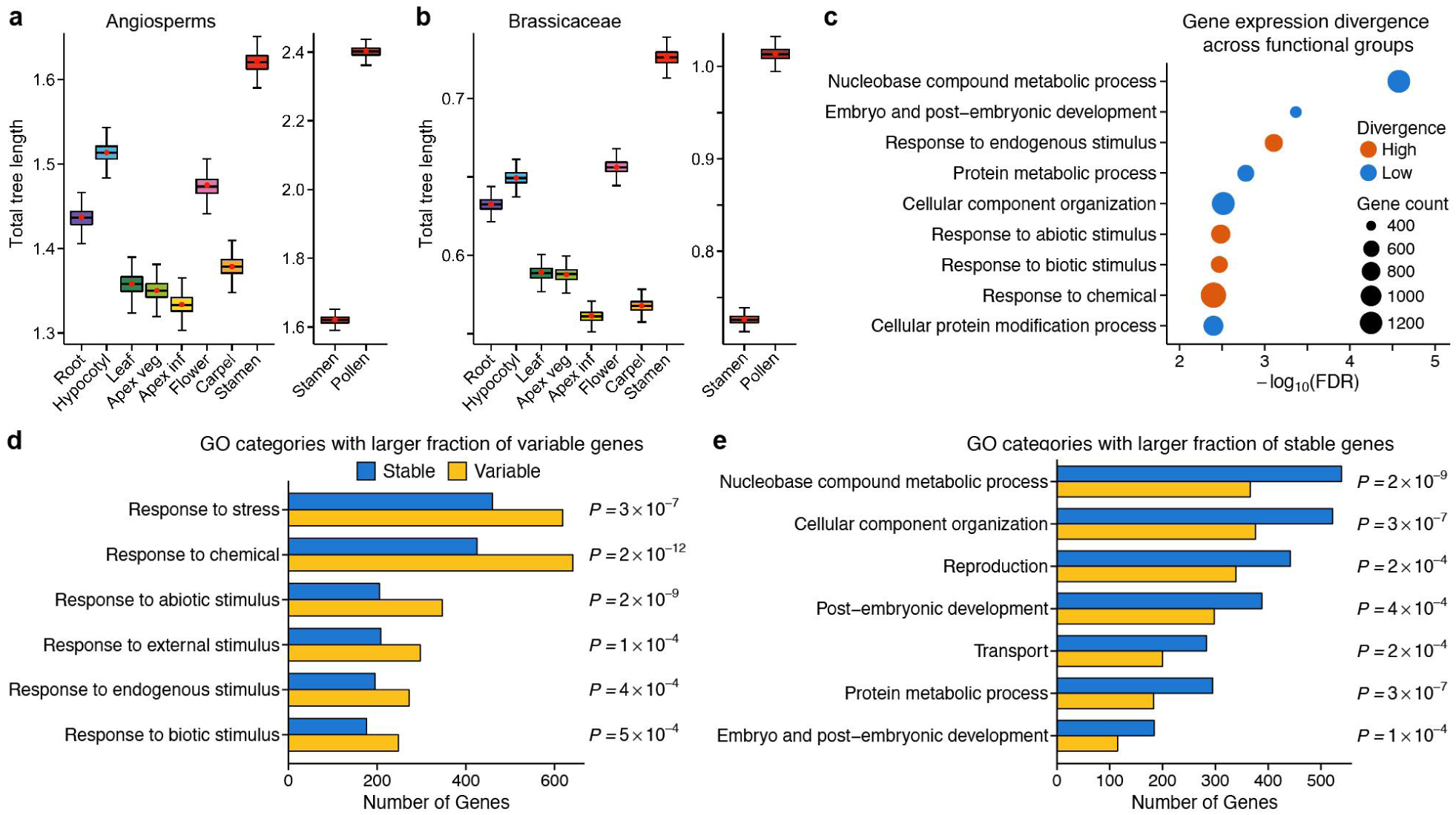
Conservation of gene expression levels among plant organs and across functional groups. **a-b,** Variations in total tree length among organs, for the angiosperm (7,003 orthologs, 7 species) (**a**) and the Brassicaceae data set (17,445 orthologs, 4 species) (**b**). Boxplots show the distribution of the total tree length from 1,000 bootstrap phylogenies; red points mark tree length of expression trees derived from unsampled expression data; whiskers are based on 1.5 times interquartile range (IQR) value. **c,** Subsets of functionally related 1-1 orthologous genes that show a higher or lower expression divergence than expression-matched control gene sets (n > 412 genes; False Discovery Rate [FDR] adjusted p-value < 0.01; two-sided Wilcoxon rank-sum test). **d-e,** Gene ontology categories that contain a larger fraction of variable (**d**) or stable (**e**) genes than expected by chance (n > 412 genes; FDR adjusted p-value < 0.001; Pearson’s chi-squared test).

We also analysed the evolutionary changes in lncRNA expression patterns. Earlier studies in both plants and animals have shown that the level of sequence conservation of lncRNAs is very low ^27, 28, 29^, and consistently, we detected only 8 orthologous non-coding transcripts across all seven angiosperm species. Therefore, we limited our analysis to younger lncRNAs that are shared between the four Brassicaceae species (Fig. 1b, n = 307). Here, expression conservation is considerably lower than what we observed for protein-coding genes, where organ-dominated clustering of samples is evident (Extended Data Fig. 8a-d). Furthermore, non-coding gene expression phylogenies display a similar branch length across all organs (Extended Data Fig. 8e,f), and branches are about twice as long as those from the protein-coding gene expression trees, illustrating rapid changes in non-coding gene expression levels (Extended Data Fig. 8f,g).

## Evolution of angiosperm gene expression levels among functional groups

Finally, we assessed the evolutionary stability of protein-coding gene expression patterns among sets of protein-coding genes that share the same biological functionality. Considering that correlation-based distances can be inaccurate when calculated from small sample sizes, we used a Monte-Carlo simulation to estimate the minimum number of samples required to obtain stable correlations ^30^. We found that subsets of expression data should contain more than 400 genes to approach the true value ρ of the population (Extended Data Fig. 9a). Therefore, we focussed on the analysis of high-level summaries of gene ontology (GO) terms and excluded all GO subsets of insufficient sample size.

As noticed earlier (Fig. 2c,d), genes that evolve slowly at the protein sequence level are on average more highly expressed than non-conserved genes. To test if a similar inverse relationship exists between gene expression level and rate of expression divergence, we divided the angiosperm ortholog expression data into sets of 500 genes, resulting in 14 quantiles of intensity distributions, and calculated the rate of expression evolution for each group. We found that quantiles containing the lowest and highest expressed genes show a lower rate of expression divergence than the rest (Extended Data Fig. 9b,c). Most of the genes enriched in both groups are involved in metabolic processes (Extended Data Fig. 9e,g). Notably, not only gene expression patterns of globally lowly or highly expressed genes are particularly conserved, but also local minima of expression levels. For example, genes that are strongly expressed in leaf or during flower development show an evolutionary conserved local minimum in root across all species (Extended Data Fig. 9d,f).

Next, we explored whether the level of gene expression evolution varies among different groups of functionally related genes. Since the above results indicated that the evolutionary stability of the expression pattern of a gene is partially related to its expression intensity, we decided to match, based on expression level and relying on numerical balance measures, two or more control groups to each GO subset of interest. We then calculated Pearson distances from pairwise species comparisons and estimated the rate of expression evolution using a non-linear regression model for both GO subset and control groups (Extended Data Fig. 9i). We observed that genes involved in the response to endogenous and external stimuli exhibited higher rates of expression divergence than controls, whereas the expression divergence of genes functioning in key cellular, metabolic and developmental processes is low (FDR-adjusted Ρ < 10^-2^, two-sided Wilcoxon rank-sum test; Fig. 4c and Extended Data Fig. 9h). In order to validate these results we calculated the coefficient of variation (CV), defined as intra-organ inter-species standard deviation normalised by the mean expression across species, for each gene as a second metric to estimate the evolutionary stability among functional groups ^31^. The outcome was very similar: The vast majority of genes that regulate metabolic, developmental and cellular processes were classified as evolutionary stable, and those that respond to abiotic, biotic, external and endogenous stimuli were primarily classified as variable (Fig. 4d,e and Extended Data Fig. 10c-e).

## Discussion

How flowering plants have evolved so rapidly and have gained ecological dominance on earth is a fundamental question in plant evolutionary biology. Numerous hypotheses have been given to explain the “mysterious”, highly accelerated rate of angiosperm diversification, including the coevolutionary interactions between pollinating insects and angiosperms, biotic seed dispersers, asexual/sexual reproduction, annual and biennial growth forms, repeated genomic rearrangements, epigenetic versatility, and “resistance” to extinction ^2, 32, 33, 34^. This study sheds light on another, angiosperm-diversity promoting component. By generating and analysing a comprehensive total RNA-seq organ transcriptome data set from seven angiosperm species, we show that angiosperm gene expression patterns evolved rapidly. Since flowering plants are sessile, exothermic organisms, they constantly have to adjust to their environment to maintain optimal growth. Our finding that genes which function in response to endogenous/external stimuli and biotic/abiotic stresses show the highest rate of expression evolution could provide one possible explanation how plants might facilitate adaptive evolution and specification.

Recent studies on angiosperm phylogenomics provide molecular evidence for two surges in angiosperm diversification. An early burst, caused by drivers such as the global change of biomes, continental rearrangements, and changes in key environmental parameters like global temperatures, occurred during the late Cretaceous Period (125-80 Myr ago). It gave rise to the majority of extant angiosperm orders and was accompanied by a substantial decline of gymnosperm plant lineages. Then, diversification proceeded at stable levels until global climatic cooling during the Cenozoic Era led to a second burst of angiosperm diversification ^35, 36, 37^. It will be interesting to link these historic events of global climate change to specific rates of gene expression evolution in future studies. This will require denser species sampling to obtain a finer resolution of gene expression divergence, and direct studies on environmental responses in specific angiosperm lineages.

## Methods

### Plant Material and conditions

All plants were grown on soil, except those used for the collection of early-stage root samples, which were grown on 0.8% plant agar plates, pH 5.8, containing ½ MS and vitamins (Duchefa M0222).

*Arabidopsis thaliana* (Col-0) was grown at 21°C in constant light with a light intensity of 170 µmoles m^-2^ s^-1^ and 65% humidity. Seeds for plants grown on soil were surface-sterilised with 70% ethanol for 5 min followed by 7% bleach + 0.1% Triton X-100 and washed three times in sterile water. Seeds were placed in water containing 0.01% agar and kept for 3 days in a 4°C room for stratification before sowing out on soil (Levington F2). Seeds for plants grown on ½ MS agar plates were surface-sterilised in a laminar flow hood with 70% ethanol + 0.1% Triton X-100 for 5 min, briefly rinsed in 96% ethanol for 1 min, air-dried and placed on plates, stratified for 3 days at 4°C and transferred to a constant light growth room.

*Arabidopsis lyrata* (MN47) ^38^, *Capsella rubella* (Monte Gargano) ^39^ and *Eutrema salsugineum* (Shandong) ^40^ were grown under the same condition described above, with the following modifications: *Arabidopsis lyrata* seeds were stratified for additional 2 weeks at 4°C after sowing out on soil. For the collection of reproductive tissues, seedlings of all three species were first grown for 2 months in short days (8h light at 21°C/16h dark at 17°C, 170 µmoles m^-2^ s^-1^ and 65% humidity) for the formation of large rosettes. Plants were then transferred to 4°C for 6-8 weeks for vernalization, and then placed in constant light to induce flowering.

*Tarenaya hassleriana* (ES1100) ^41^ was germinated in Parafilm-sealed petri dishes on filter paper soaked with water in long days (16h light at 28°C/ 8h dark at at 20°C) with a light intensity of 300 µmoles m^-2^ s^-1^ and 65% humidity. Seedlings were grown on soil in constant light (22°C, 300 µmoles m^-2^ s^-1^, 65% humidity). After a few weeks, plants were transferred into larger pots and light levels were increased to 400 µmoles m^-2^ s^-1^. For root collections, seeds were germinated and seedlings were grown on filter paper soaked with water containing ½ MS.

*Medicago truncatula* (A17) ^42^ seeds were sterilised for 5 min in concentrated H_2_SO_4_, washed three times in sterile water, treated 3 min with 7% bleach, washed again three times in sterile water, and soaked in sterile water for 2-3 hours. Seeds were transferred to ½ MS plant agar plates (pH 5.8), which were sealed with Parafilm, wrapped with aluminium foil and stored upside-down at 4°C for 2-3 days. Plates were transferred to 20°C and kept in darkness upside-down for 1 day. Seedlings were kept on plates (for root sample collections), or transferred into large pots containing soil (Levington M3) and grown in constant light at 21°C with a light level of 350 µmoles m^-2^ s^-1^ and 60% humidity.

*Brachypodium distachyon* (Bd21) ^43^ seeds were planted into soil and stratified for one week in a 4°C room to synchronise germination. Plants were grown in constant light at 23°C with a light level of 220 µmoles m^-2^ s^-1^ and 65% humidity. For plant culture, the palea and lemma were removed with forceps under a microscope. Seeds were sterilised in 7% bleach with 0.1% Triton X-100 for 5 min, rinsed three times with sterile water, incubated for 1 min in 96% ethanol, and rinsed again 3 times in sterile water. For mesocotyl collections, seeds were planted at least 2 cm deep into the soil.

### Sample collection and RNA-Seq library preparation

All samples were collected in 1.5 ml Safe-Lock microcentrifuge tubes floating on liquid nitrogen, except larger-sized organs, which were collected in 50 ml centrifuge tubes on dry ice. Depending on sample type, between 25 and 100 organs were pooled as one replicate. All samples were collected in biological triplicates.

Full details on organ dissections can be found in Supplementary Table 1. For the comparative samples, we used *Arabidopsis thaliana* as a reference model and sampled the organs of the other species accordingly (Supplementary Table 1). The following organ developmental stages of *Arabidopsis thaliana* were used as reference: (1) Roots from 5 day old plants, dissected at approximately 3 mm from the apex; (2) hypocotyl from 10 day old plants; (3) leaves 1+2 from 7 day old plants; (4) dissected vegetative meristem from 7 day old plants; (5) dissected inflorescence meristem from 21 day old plants. All flowers older than stage 4/5 were removed ^44^; (6) flower stage 12 ^44^; carpel (7) and stamen (8) from flower stage 12 ^44^; (9) mature pollen. Roots, meristems and floral organs were dissected using forceps under a microscope. Mature pollen was collected as described previously ^45^.

Dissected organs were homogenised and total RNA was isolated using RNeasy Plant Mini Kit (QIAGEN 74904), following the manufacturer’s instructions. RNA integrity was measured using an Agilent 2200 TapeStation. Strand-specific total RNA-Sequencing libraries were prepared using the TruSeq Stranded Total RNA with Ribo-Zero Plant kit according to the manufacturer’s specifications (Illumina 20020610). ERCC RNA Spike-In Control Mix (Ambion 4456740) was added at the beginning of the library preparation before the ribosomal RNA (rRNA) depletion step to allow sample-specific estimation of the lower detection limit. The quality of the generated libraries was assessed using an Agilent 2200 TapeStation. All RNA-Seq libraries were sequenced on the Illumina HiSeq 4000 system by BGI Beijing, except five libraries that were sequenced in-house on the Illumina NextSeq 500 platform, producing paired-end 75bp reads. Depending on organ type, libraries were sequenced to a depth of 50M to 200M reads per sample to obtain an equal amount of mapped, deduplicated reads (Extended Data Fig. 1a-c). This was necessary due to incomplete removal of cytoplasmic, mitochondrial and chloroplast rRNAs especially in leaf samples. To rule out potential biases during the library preparation step, we generated technical replicates for some of the samples. We also re-sequenced samples on different flow cells, which did not reveal any obvious biases during the sequencing step.

### Read mapping, transcript annotation and orthology assignment

Genomic annotations were retrieved from Phytozome ^46^, v12 (for *A. lyrata*, *C. rubella*, *E. salsugineum*, *M. truncatula*, and *B. distachyon*), Ensembl ^47^, release 34 (for *A. thaliana*) and release 92 (for *H. sapiens*, *P. troglodytes*, *P. paniscus*, *G. gorilla*, *P. pygmaeus*, *M. mulatta*, *M. musculus*, and *M. domestica*), and RefSeq ^48^ (for *T. hassleriana*). We expanded the angiosperm annotations to include novel protein-coding genes, splice junctions and lncRNAs as described ^49^, Briefly, we used STAR ^50^ to align deduplicated reads to the reference genome, Cufflinks ^51^ for transcriptome assembly, GeMoMa ^52^ to calculate splice junction coverage, and TransDecoder ^53^ to identify candidate coding regions within transcripts with unknown coding potential.

Using *A. thaliana* as a reference species for plants and *H. sapiens* as reference for animals, we performed proteome-wide orthology inference using orthologr ^54^. Here, pairwise 1-1 orthologous protein-coding genes between *A. thaliana*/*H. sapiens* and any of their respective subject species were retrieved using BLASTP best reciprocal hits of the longest splice variant using an e-value threshold of 1E-5, query-length-coverage threshold of 70%, and sequence identity threshold of 30%. For the detection of 1-1 orthologous non-coding RNAs, we also performed best reciprocal BLASTP hits using an e-value threshold of 1E-2, query-length-coverage threshold of 70%, and sequence identity threshold of 30%. Computationally reproducible analysis scripts can be found at https://github.com/HajkD/DevSeq.

### Expression level estimation, threshold, data normalisation, and biological data analysis

After mapping the clean raw reads to the reference genome with STAR and generating raw read counts, we normalised the count data using DESeq2 ^55^. We calculated expression levels as transcript per million (TPM), and transformed the normalised counts using variance stabilizing transformation (VST) implemented in DESeq2. VST counts were used as input for hierarchical clustering (hclust) and Principal Component Analysis (PCA). For most other analyses, we used TPM values as input.

Biological data analysis was done in R ^56^, using plyr ^57^, dplyr ^58^, and data.table ^59^ for data manipulation, R.utils ^60^ and devtools ^61^ for development and utility functions, matrixStats ^62^ for optimised functions that apply to matrices, and ggplot2 ^63^, gplots ^64^, scales ^65^, gtable ^66^, grid ^56^ and ggbeeswarm ^67^ for creating graphics, graphics layout and plotting. To retrieve the number of expressed genes in each sample, we applied a threshold based on ERCC spike-ins. We applied the 0.05 percentile of detected spike-ins throughout the study. A gene was considered as expressed if its expression value was above this threshold level in at least two out of three biological replicates.

Hierarchical clustering of pairwise distances, preserved as metric ^68^, was done using the ‘hclust’ and ‘dendrapply’ functions of the ‘stats’ package in R. Dendrogram objects were extended using the R packages dendextend ^69^ and factoextra ^70^. Principal Component Analysis was done using the ‘prcomp’ function of the ‘stats’ package.

To retrieve pairs of adjacent transcripts, we used the Bioconductor ^71^ packages GenomicRanges ^72^ and rtracklayer ^73^. We categorised gene pairs as overlapping cisNAT/protein-coding gene pair (NAT/PC), overlapping protein-coding/protein-coding gene pair (PC/PC), and non-overlapping, neighbouring protein-coding gene pair of the same (PCSS) or opposite (PCOS) strand. Only genes with an expression value of 0.5 TPM were considered.

Neighbour-joining gene expression phylogenies were constructed using the ape ^74^ and ggtree ^75^ packages. VST count data of either core orthologs (n = 7003) or Brassicaceae orthologs (n = 17,445) was used as input, and the pairwise distances between samples were calculated as Pearson distances, preserved as metric. The accuracy of branching patterns was estimated by random sampling with replacement (n = 1,000 bootstraps), and tree nodes were colour-coded according to the results of the bootstrap analysis.

### Rate of expression evolution

To estimate the rate of expression evolution for each organ, we first calculated pairwise intra-organ Pearson distances as metric between all species pairs across the entire phylogenetic tree. The distances were assigned to their corresponding estimated species divergence time, retrieved from the TimeTree of Life ^11^. Next, we fitted a negative exponential growth model,

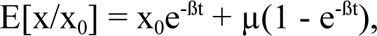

where E[x/x_0_] is the mean, ß is the rate of expression evolution, t is evolutionary time, and µ is the optimal expression value, using the ‘nls’ function of the ‘stats’ package in R. This non-linear model assumes that gene expression divergence is predominantly under stabilising selection and saturates with evolutionary time. We also applied locally estimated scatterplot smoothing (LOESS) as a non-parametric method. We then determined the average slope value of the regression fits for each individual organ as a measure of transcriptome evolution.

As a second measure, we estimated the transcriptome distance between species based on the Ornstein-Uhlenbeck (OU) model with variable optimal expression level among genes, implemented in the R package TreeExp2 ^76^. For the analysis of the mammalian data set ^3^, biological replicates that showed a sample correlation below 0.85 (Pearson’s r) were excluded.

### Gene expression evolution among groups of functionally related genes

Sample correlations converge to the true value ρ of the population with increasing sample size, and it has been shown, using Monte-Carlo simulations to determine the point of stability (POS), that sample size should approach n = 250 for stable estimates ^30^. We adapted this approach to our ortholog expression data set, using two transcriptome samples, *Brachypodium distachyon* carpel and *Tarenaya hassleriana* carpel, with an actual correlation of ρ = 0.6 of the population. We drew 1,000 bootstrap samples from this population, and for each bootstrap sample we calculated the correlation for every sample size, starting with n = 20 to a maximum sample size of n = 1,000, with a step size of 1. Based on these 1,000 trajectories of correlation, we calculated the POS, defined as the critical sample size from which at least 80% of all trajectories do not leave the corridor of stability (w = 0.1) around the true value ρ anymore. Based on the results of this simulation, we limited the analysis to high-level summaries of gene ontology (GO slim) terms with a sample size of n > 412 genes. GO slim categories were retrieved from The Arabidopsis Information Resource (TAIR) (https://www.arabidopsis.org).

Because the evolutionary stability of different gene sets seems to be partially related to their average expression level ^31^ (Extended Data Fig. 9b,c), we decided to match, based on expression level and relying on numerical balance measures ^77^, two or more control groups to each GO subset of interest. To find the optimal number of matched samples (control groups), we focussed on two balance measures: the standardised mean difference between treatment and control means, with a target value of ≤ 0.01, and the variance ratio between treatment and control, with a target value of ≥ 0.99. We performed nearest distance matching using the ‘MatchIt’ package ^78^ in R. For expression-matching the groups of stable and variable orthologous genes (Extended Data Fig. 10c,d), we performed nearest distance matching with the caliper option and a 1:1 control to treatment ratio. The rate of transcriptome evolution for the GO subsets and the control groups was calculated as described above.

### Web platform for plant evolutionary transcriptomics

We developed an interactive web application for data analysis and visualisation, based on the LAMP (Linux, Apache, MySQL, PHP/Python) software stack for server-side database management and programming. The application uses the ‘pandas’ library ^79^ for data analysis and manipulation, and the ‘SciPy’ library ^80^ for hierarchical clustering in Python. Client-side data visualisation is implemented in JavaScript using the D3.js ^81^, c3.js ^82^ and Plotly.js ^83^ chart libraries and custom scripts. The web platform can be accessed at https://www.devseqplant.org and the source code can be found at https://github.com/schustischuster/DevSeqPlant.

## Acknowledgements

We thank Detlef Weigel, Aram Gurzadyan, Tyler Gibson and Eldad Afik for valuable scientific discussions, and Sandra Cortijo for advice on RNA-seq library preparation. For financial support we thank the Gatsby Charitable Foundation (GAT3731/DAA to E.M.M. and GAT3272C to O.L.) and the German Research Foundation (DFG INST 271/339-1 FUGG, FZT 118, GR 3526/2, GR 3526/6, GR 3526/7, and GR 3526/8 to I.G. The E.M.M. lab is also supported by the Howard Hughes Medical Institute. H.G.D. was partly supported by the Max Planck Society. A.G. and H.G.D. thank the Stickstoffwerke Piesteritz for the SKWP Research Prize 2013 and 2017.

This article is subject to the Howard Hughes Medical Institute’s Open Access to Publications policy. HHMI lab heads have previously granted a nonexclusive CC BY 4.0 license to the public and a sublicensable license to HHMI in their research articles. Pursuant to those licenses, the author-accepted manuscript of this article can be made freely available under a CC BY 4.0 license immediately upon publication.

---

**Extended Data Fig. 1.**
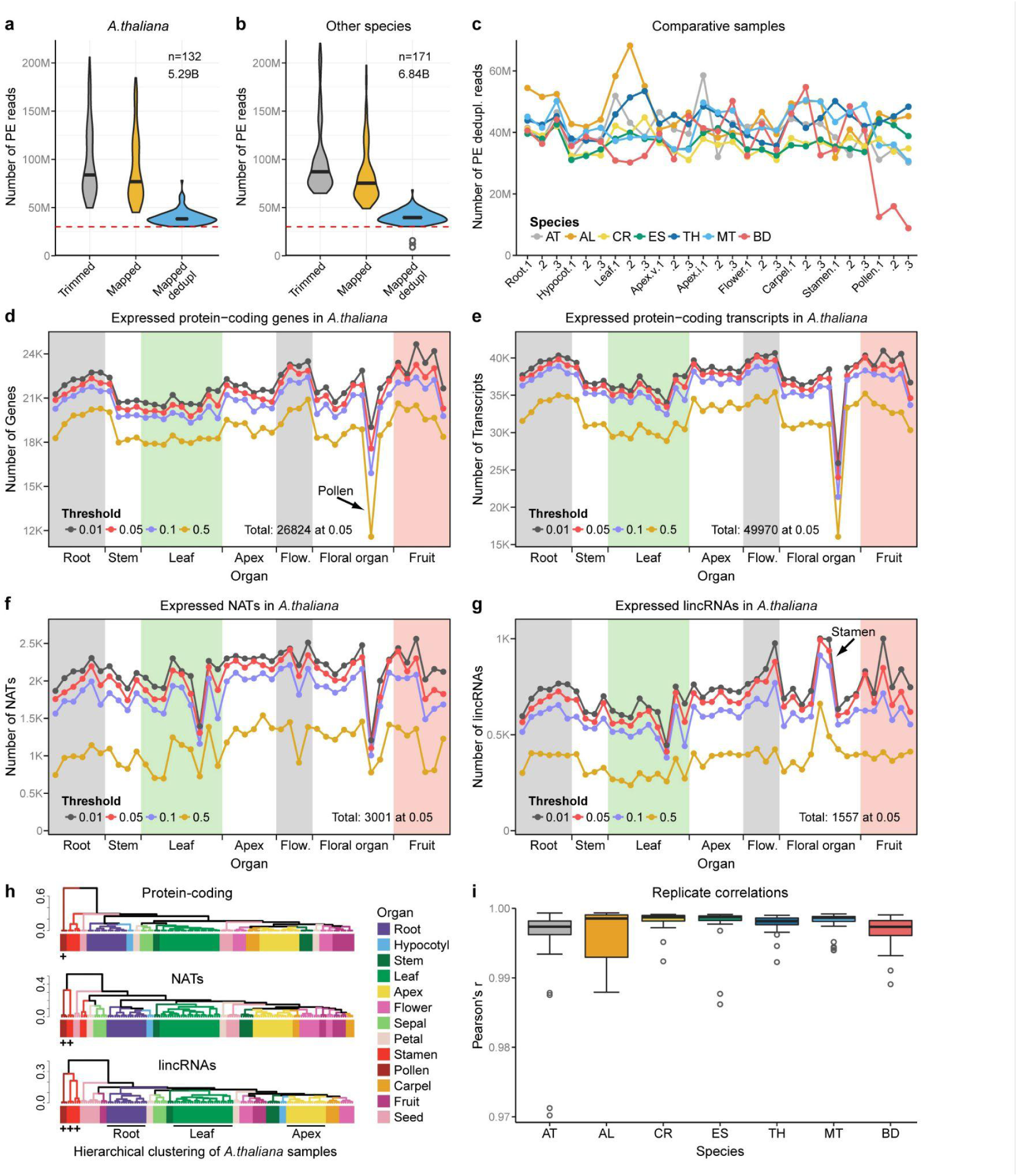
Transcriptome dynamics throughout *Arabidopsis thaliana* development. **a,** Number of paired-end trimmed, mapped and mapped/deduplicated reads in *Arabidopsis thaliana* (n = 132 samples) (**a**), and in the other plant species (n = 171) (**b**). **c,** Number of paired-end mapped, deduplicated reads in the comparative samples. AT, *Arabidopsis thaliana*; AL, *Arabidopsis lyrata*; CR, *Capsella rubella*; ES, *Eutrema salsugineum*; TH, *Tarenaya hassleriana*; MT, *Medicago truncatula*; BD, *Brachypodium distachyon*. Hypocotyl tissue was replaced by mesocotyl in *Brachypodium*. **d-g,** Number of detected protein-coding genes (**d**), protein-coding transcripts (**e**), cis-natural antisense transcripts (NATs) (**f**), and long intergenic non-coding RNAs (lincRNAs) (**g**) throughout AT development. ERCC spike-ins were used to determine the lower limit of detection, with 0.01, 0.05, 0.1 cutoff corresponding to quantiles of ERCC expression level; 0.5, fixed threshold of 0.5 TPM. Absolute numbers of expressed genes are supported by expression in at least two of three biological replicates in one or more organs and species above threshold. **h,** Hierarchical clustering of pairwise Pearson distances for AT samples based on protein-coding and non-coding gene expression using average linkage method. Samples are colour-coded according to the organ. Crosses denote the main outgroup. **i,** Distribution of Pearson correlations of pairwise comparisons of all three biological replicates across samples.

**Extended Data Fig. 2.**
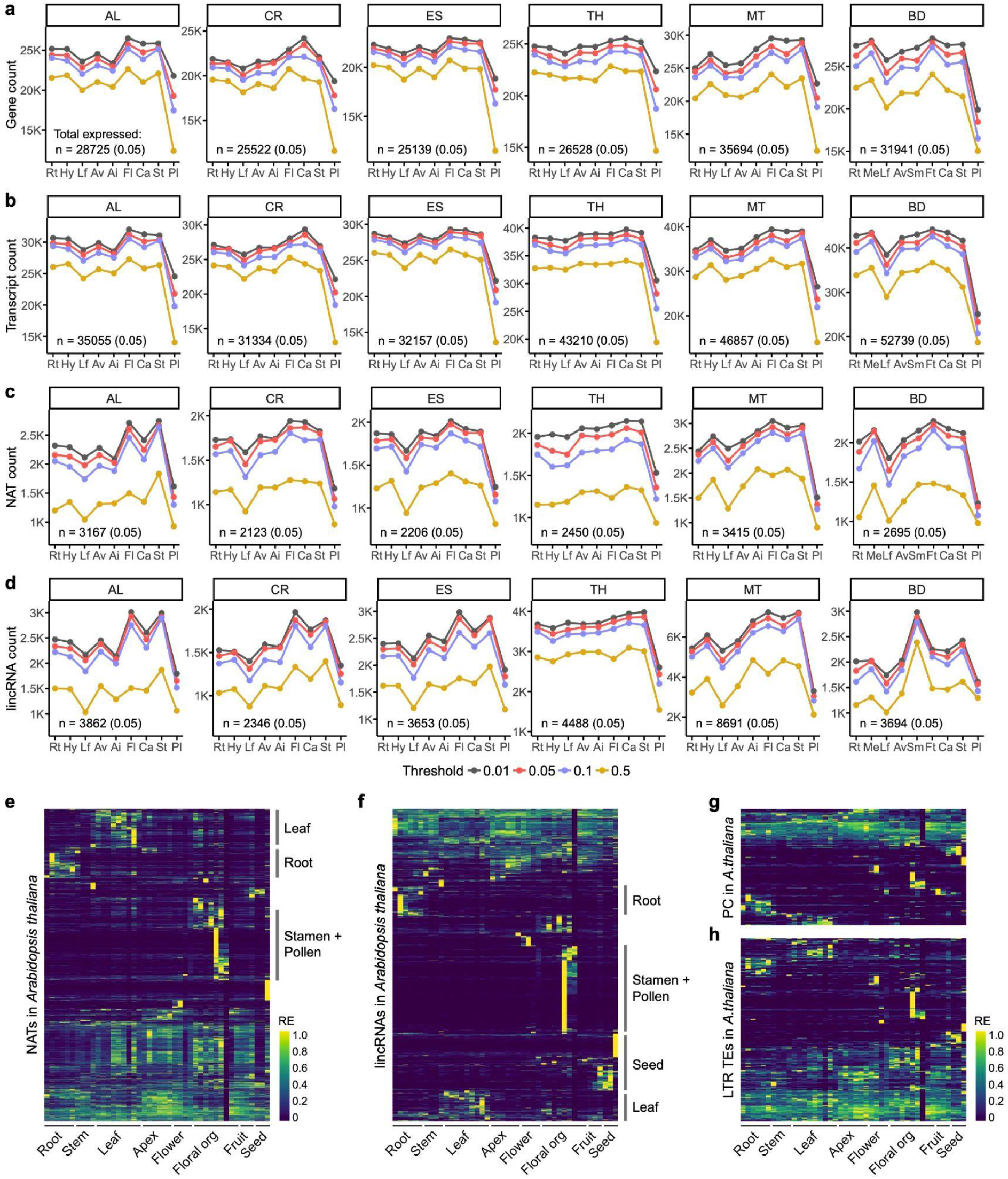
Coding and non-coding transcriptome dynamics across angiosperm organs. **a-d,** Number of detected protein-coding genes (**a**), protein-coding transcripts (**b**), cis-natural antisense transcripts (NATs) (**c**), and long intergenic non-coding RNAs (lincRNAs) (**d**) across plant organs and species. Rt, root; Hy, hypocotyl; Lf, leaf; Av, vegetative apex; Ai, inflorescence apex; Fl, flower; Ca, carpel; St, stamen; Pl, pollen; Me, mesocotyl; Sm, lateral spikelet meristem; Ft, floret; AT, *Arabidopsis thaliana*; AL, *Arabidopsis lyrata*; CR, *Capsella rubella*; ES, *Eutrema salsugineum*; TH, *Tarenaya hassleriana*; MT, *Medicago truncatula*; BD, *Brachypodium distachyon*. ERCC spike-ins were used to determine the lower limit of detection, with 0.01, 0.05 and 0.1 cutoff corresponding to quantiles of ERCC expression level; 0.5 represents a fixed threshold of 0.5 TPM. Absolute numbers of expressed genes are supported by expression in at least two of three biological replicates in one or more organs and species above threshold. **e-h,** Expression pattern of NATs (**e**), lincRNAs (**f**), newly annotated protein-coding genes (**g**) and active LTR retrotransposons (**h**) throughout *Arabidopsis thaliana* development. Organ-specific expression of long non-coding RNAs is particularly evident in pollen, stamen and seed. Expression estimates (TPM) were scaled to the unit interval [0, 1], and genes were hierarchically clustered. Plots were generated using the DevSeq web application (https://www.devseqplant.org).

**Extended Data Fig. 3.**
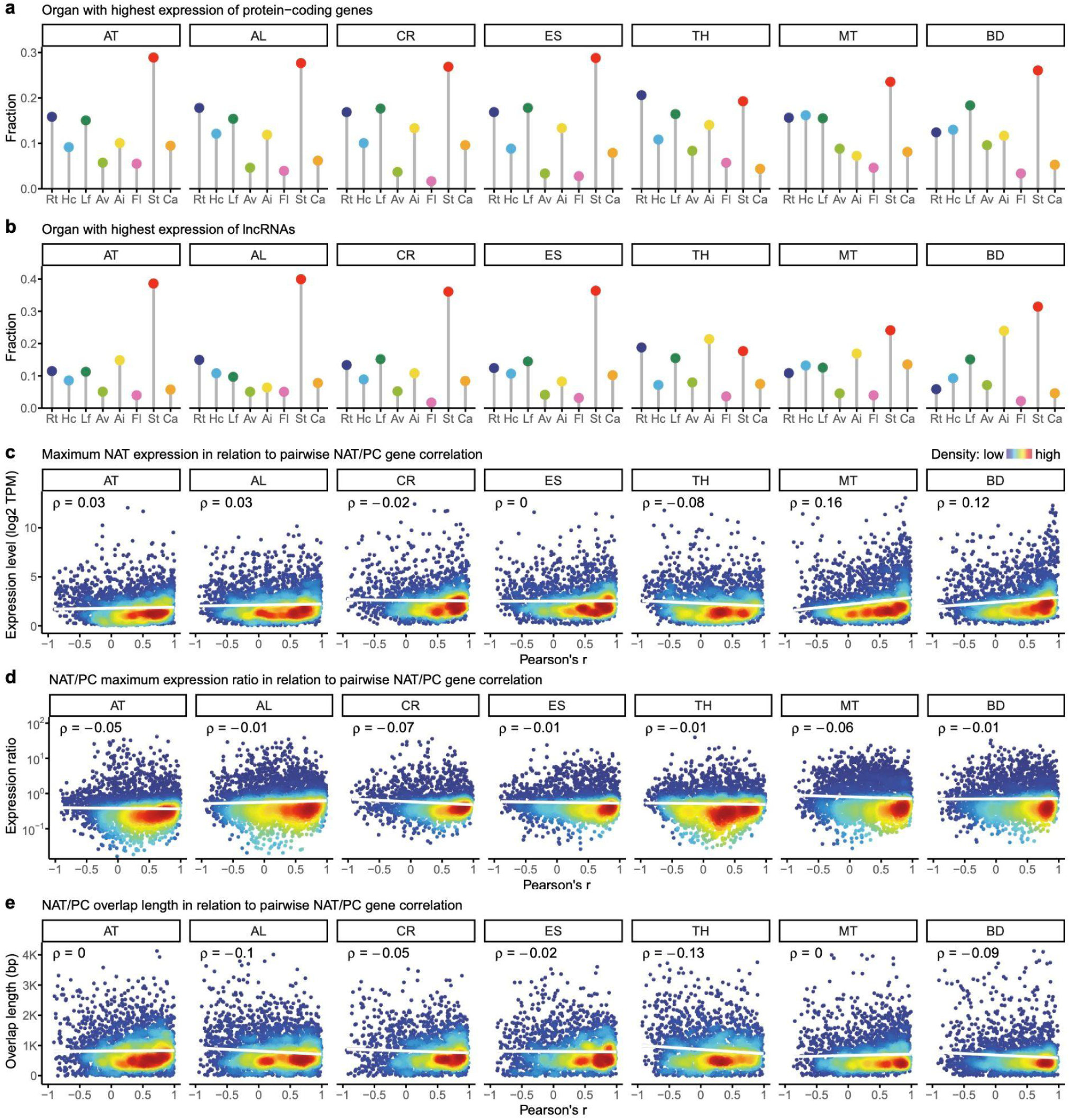
Protein-coding and lncRNA expression patterns across angiosperms. **a-b,** Fraction of protein-coding genes (**a**) and long non-coding RNAs (**b**) in which maximum expression was observed in eight organs in seven angiosperm species. Rt = root, Hc = hypocotyl, Lf = leaf, Av = Apex vegetative, Ai = Apex inflorescence, Fl = flower, St = stamen, Ca = carpel. AT = *Arabidopsis thaliana*, AL = *Arabidopsis lyrata*, CR = *Capsella rubella*, ES = *Eutrema salsugineum*, TH = *Tarenaya hassleriana*, MT = *Medicago truncatula*, BD = *Brachypodium distachyon*. **c-e**, Relationship between the co-expression of cis-natural antisense transcript (cis-NAT)/protein-coding gene (PC) pairs, calculated as pairwise Pearson correlation coefficient, and the maximum expression level of cis-NATs (**e**), the expression ratio of cis-NAT/PC transcript pairs (**f**), and cis-NAT/PC overlap length (**e**).

**Extended Data Fig. 4.**
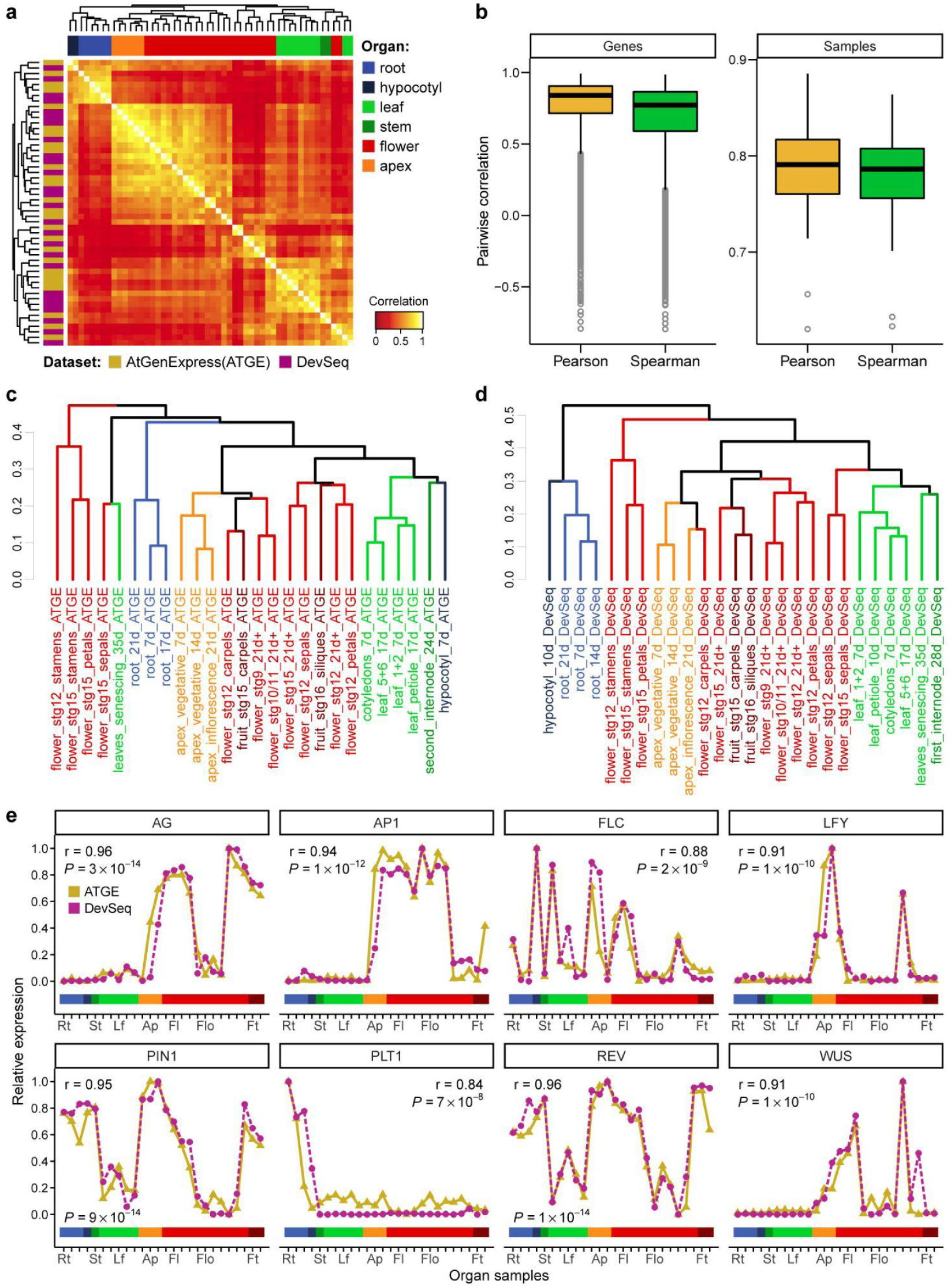
Qualitative analysis between DevSeq and AtGenExpress data sets. **a,** Symmetrical heat map of Pearson correlations from AtGenExpress genome array and DevSeq RNA-Seq gene expression profiles. Each data set consists of mRNA expression profiles from 19243 genes across 26 comparative samples. Expression estimates (gcRMA and TPM for AtGenExpress and DevSeq, respectively) were linearized and scaled to the unit interval [0, 1] to adjust for differences in the dynamic range. Samples were clustered based on Pearson distance using complete linkage method. **b,** Correlation coefficients from pairwise comparisons of gene or sample profiles from the AtGenExpress and DevSeq datasets. Data was normalised and scaled as in (a). **c-d**, Hierarchical clustering dendrograms of AtGenExpress (c) and DevSeq (d) samples based on Pearson distance using complete linkage analysis. Differences are found for the clustering of hypocotyl, fruit, and senescing leaf. **e,** Gene expression profiles of selected marker genes in AtGenExpress and DevSeq across organs and developmental stages. Expression estimates were log-transformed and scaled to the unit interval [0, 1]. Pearson correlations are shown in each plot. Organ colours are shown as in (a).

**Extended Data Fig. 5.**
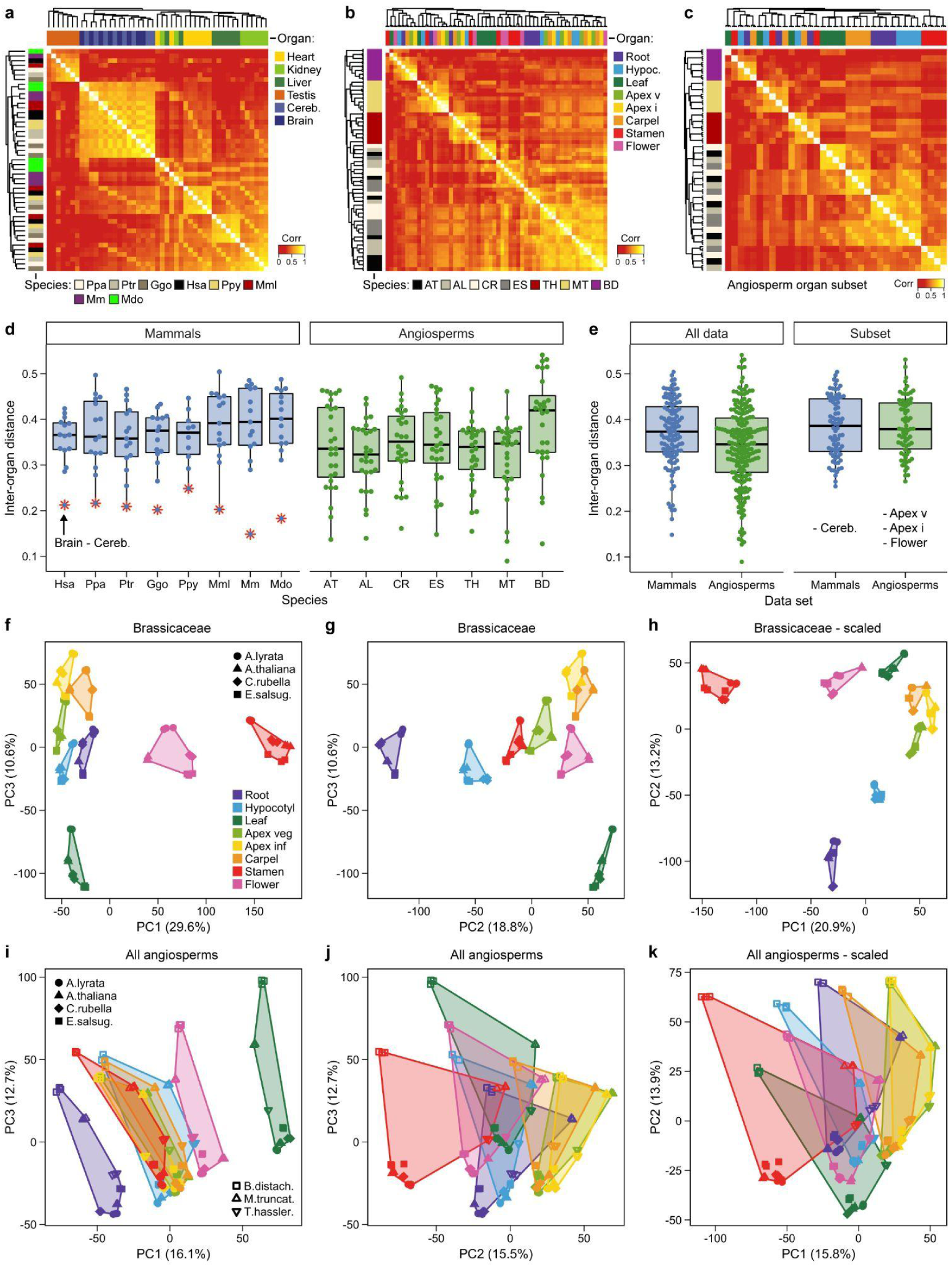
Conservation of protein-coding gene expression in angiosperms and mammals. **a,** Symmetrical heatmap of Pearson correlations from mammalian organ transcriptomes. Expression data from Brawand et al. {Brawand, 2011} has been re-analysed using our DevSeq RNA-seq pipeline. Expression levels from 7,024 1-1 orthologous protein-coding genes, estimated as normalised counts after variance-stabilising transformation (VST), were analysed across six organs and eight species. Samples were hierarchically clustered based on Pearson distances using average linkage method. Colour bars: horizontal, organs; vertical, species. **b-c,** Symmetrical heatmap of Spearman correlations from angiosperm organ transcriptomes using the complete dataset (**b**), and a subset of organs without apex tissues and flower (**c**). Expression levels of 7003 1-1 orthologous protein-coding genes were estimated as VST counts, and samples were clustered based on Spearman distances using average linkage method. **d-e,** Distribution of inter-organ Pearson distances among the mammalian and angiosperm datasets, shown for each species individually (**d**), and as combined data (**e**). Asterisks in (**d**) indicate brain-cerebellum interorgan distance. Subset in (**e**) shows the distribution of distances after removing the most similar organ pairs from the dataset. **f-k,** Principal-component analysis (PCA) of protein-coding gene expression levels from 17,445 1-1 orthologous protein-coding genes in Brassicaceae (**f-h**), and from 7,003 1-1 orthologs protein-coding genes conserved across all seven angiosperm species (**i-k**).

**Extended Data Fig. 6.**
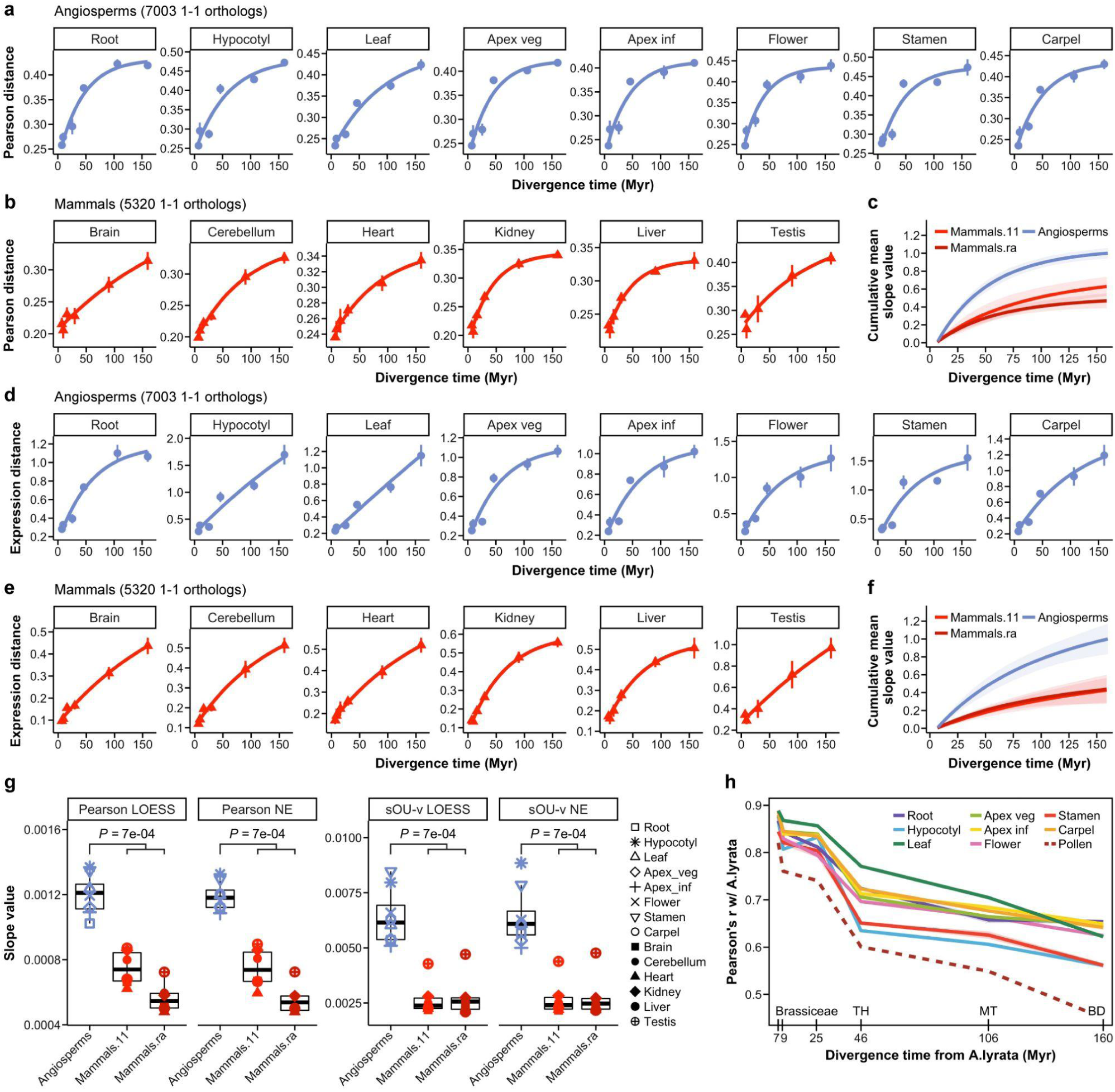
Angiosperm and mammalian transcriptome distances. **a–f,** Average pairwise expression distance across organs for angiosperm (**a, d**) and mammalian (**b, e**; data from Brawand, 2011) species, calculated as Pearson distance (**a, b**) or as expression distance estimated under the Ornstein-Uhlenbeck (OU) model (**d, e**). Each data point represents the average of the transcriptome distances from all species pairs that share the same last common ancestor. Non-linear regression fit using a negative exponential growth model is illustrated as a solid line. Error bars, standard deviation of the mean across species pairs. **c, f,** Cumulative mean slope value of expression distances, measured as Pearson distance or as distance estimated under the OU model (**f**), from angiosperm and mammalian organs. Shaded areas show 95% confidence intervals. Mammals.11, expression data published in Brawand, 2011; Mammals.ra, re-analysed data from Brawand, 2011, using our RNA-seq expression quantification and detection pipeline; Angiosperms, DevSeq angiosperm expression data. **g,** Distribution of the average slope values, derived from locally estimated scatterplot smoothing (LOESS) and negative exponential growth (NE) regression fittings, for Pearson and OU-based models across angiosperm and mammalian organs. Different point shapes denote individual organs. Organ gene expression levels diverge faster in angiosperms than in mammals (Ρ = 10^-4^, two-sided Wilcoxon rank-sum test). **h,** Pairwise Pearson correlations between *Arabidopsis lyrata* and the other species, for 8 organs and pollen, based on mRNA gene expression levels (log-transformed TPM; 7,003 genes). For Brachypodium, mesocotyl was analysed instead of hypocotyl. TH, *Tarenaya hassleriana*; MT, *Medicago truncatula*; BD, *Brachypodium distachyon*.

**Extended Data Fig. 7.**
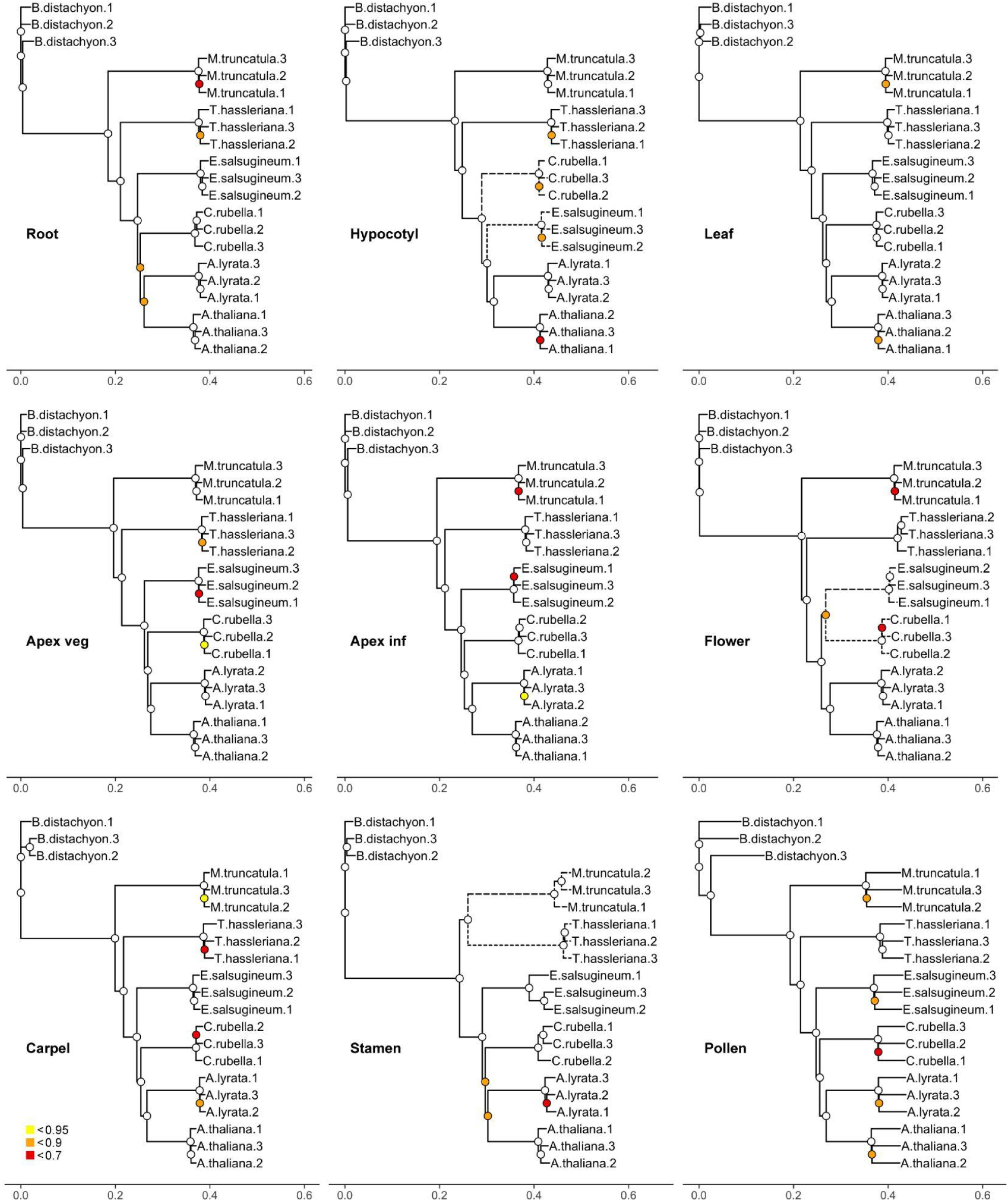
Angiosperm phylogenies based on gene expression evidence. Neighbour-joining trees based on pairwise Pearson distances, preserved as metric, for angiosperm organs. Node colours show the accuracy of branching patterns estimated by random sampling with replacement (7,003 1–1 orthologous protein-coding genes; 1,000 bootstraps). Dashed lines mark branches that differ from taxonomic relationships.

**Extended Data Fig. 8.**
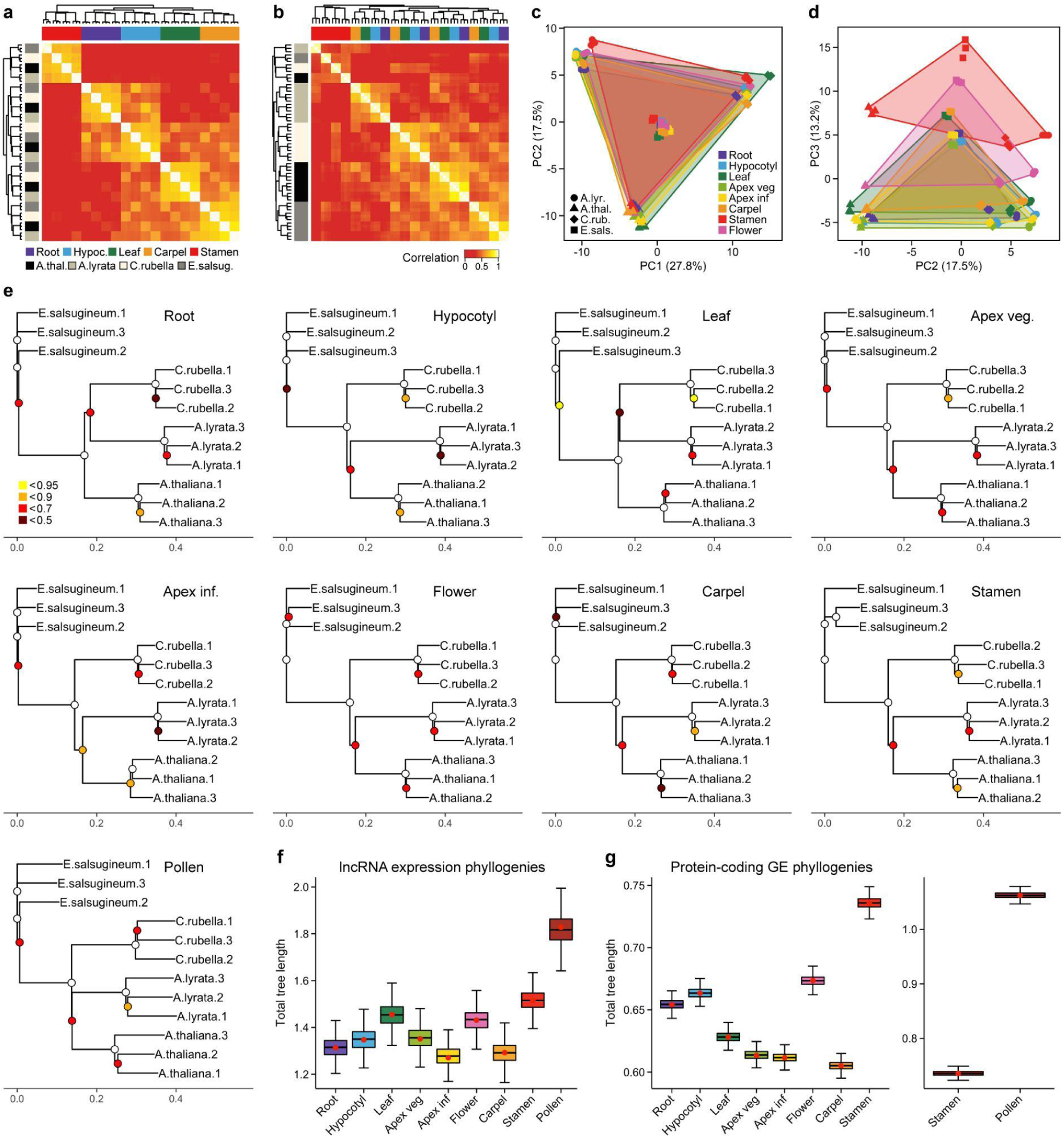
Evolution of non-coding gene expression levels in angiosperms. **a-b,** Symmetrical heatmap of Spearman correlations from Brassicaceae organ transcriptomes, for protein-coding genes (**a**, n = 17,445) and lncRNAs (**b**, n = 307). Expression levels, estimated as normalised counts after variance-stabilising transformation (VST), were analysed across the five most distant organs from four angiosperm species (*Arabidopsis thaliana*, *Arabidopsis lyrata*, *Capsella rubella*, *Eutrema salsugineum*). Samples were hierarchically clustered based on Spearman distances using average linkage method. Colour bars: horizontal, organs; vertical, species. **c-d,** Principal-component analysis (PCA) of gene expression levels from 307 lncRNA orthologs shared between the four Brassicaceae species. **e,** Neighbour-joining trees based on pairwise Spearman distances, preserved as metric, for Brassicaceae non-coding organ transcriptomes. Node colours show the accuracy of branching patterns estimated by random sampling with replacement (307 1-1 orthologous lncRNAs; 1,000 bootstraps). **f-g,** Total tree length of lncRNA (f, n = 307) and protein-coding gene (g, n = 17,445) expression trees, based on pairwise Spearman distances, from eight organs in four angiosperm species. Boxplots show the distribution of the total tree length from 1,000 bootstrap phylogenies.

**Extended Data Fig. 9.**
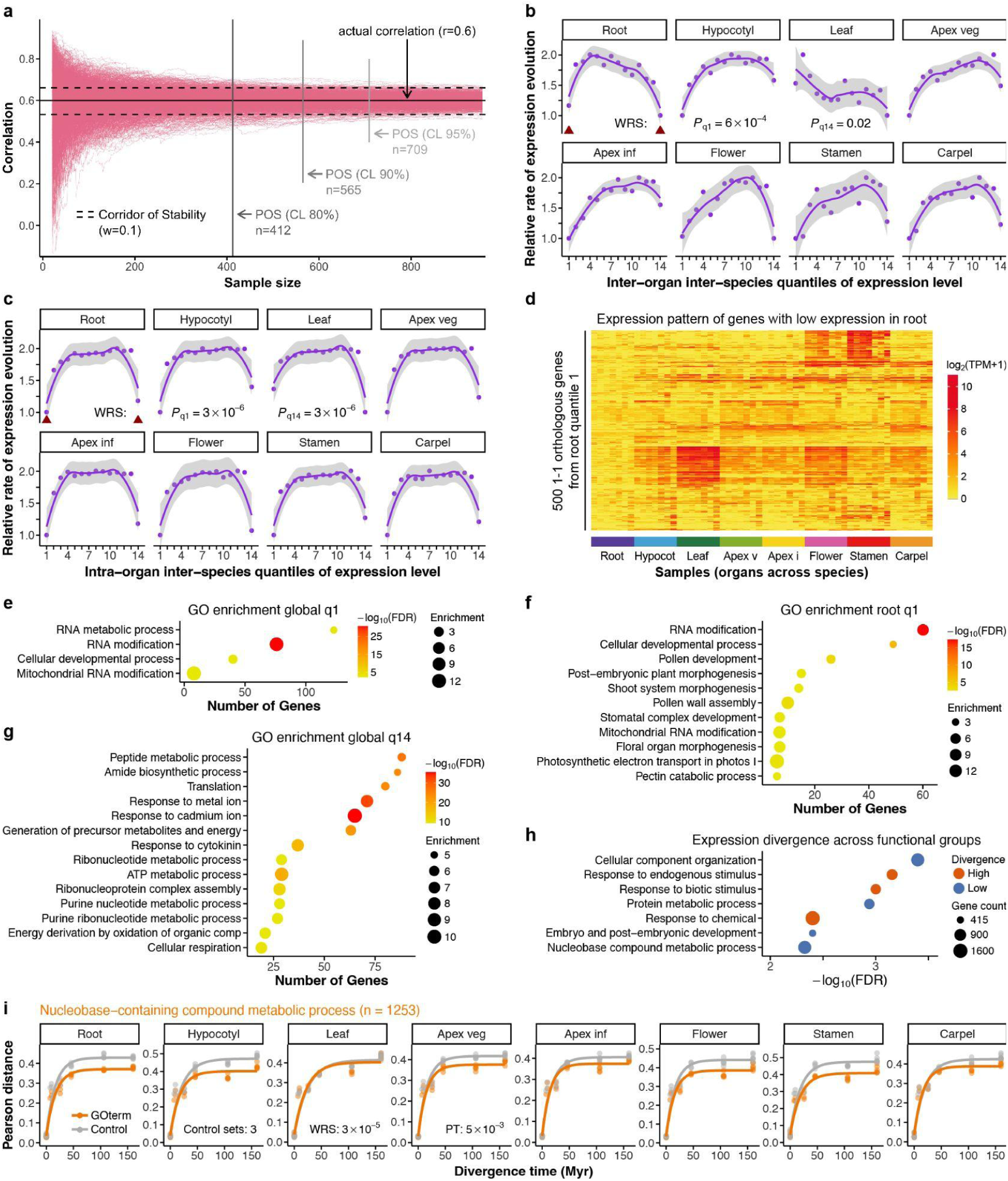
Evolution of angiosperm gene expression levels across functional groups. **a,** Monte-Carlo simulation to determine the critical sample size required to obtain stable correlations. Lines show bootstrapped trajectories of an actual Pearson correlation, with the final correlation of r = 0.6. POS, point of stability at various confidence levels (80%, 90%, 95%). **b-c,** Rate of expression evolution for inter-organ inter-species (**b**) and intra-organ inter-species (**c**) quantiles of expression level from 7,003 angiosperm 1-1 orthologous genes (14 quantiles with n = 500 genes each). Rates of expression evolution were scaled to the unit interval [1, 2]. Genes from the lowest and the highest quantile of expression level (arrow heads) show a lower expression divergence than the rest (P < 0.05, two-sided Wilcoxon rank-sum test). **d,** Expression pattern of the 500 most lowly expressed genes from root (left arrowhead in **c**) across organs and species. Samples were clustered based on pairwise Pearson distance using complete linkage method, and grouped by organ. **e-g,** High-level summaries of GO term categories identified among orthologous genes from the lowest (**e**) and highest (**g**) inter-organ inter-species quantile of expression level, and among orthologous genes expressed at very low level in root (**f**). **h,** Subsets of functionally related 1-1 orthologous genes that show a higher or lower expression divergence than expression-matched control gene sets (n > 412 genes; FDR adjusted p-value < 0.01; permutation test with 100,000 iterations). **i,** Pearson distances for genes mapping to the GO term “Nucleobase-containing compound metabolic process”, and for expression-matched control sets. Each data point represents the transcriptome distance from a species pair. Non-linear regression fit is illustrated as a solid line.

**Extended Data Fig. 10.**
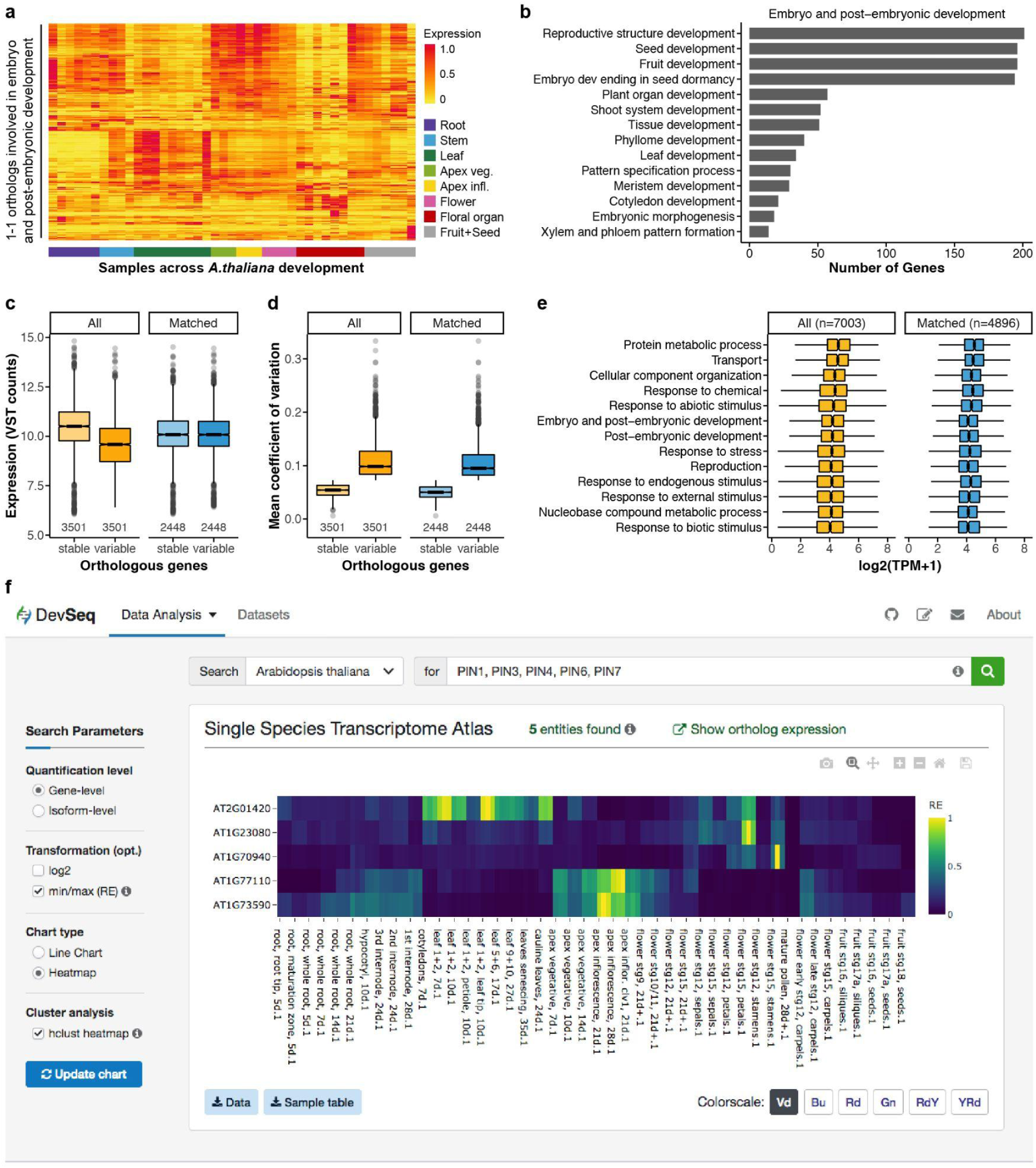
Developmental genes, gene expression variation across species, and web-based analysis platform. **a,** Expression pattern of 1-1 orthologous protein-coding genes involved in embryo and post-embryonic development in *Arabidopsis thaliana* (n = 415). Expression estimates were scaled to the unit interval [0, 1]. Samples were hierarchically clustered based on Pearson distances, using average linkage method, and grouped by organ and developmental stage. **b,** Gene ontology (GO) terms present in the GO subset “Embryo and post-embryonic development”. **c-d,** Distribution of expression values (**c**) and mean coefficient of variation (standard deviation across species normalised by mean expression, **d**) among 1-1 orthologous protein-coding genes that were categorised as stable or variable expressed across species, for all genes (n = 3501 genes/category), and for expression-matched genes (n = 2448 genes(category). **e,** Distribution of expression values among groups of functionally related genes (GO subsets), for all (n = 7003) and for expression-matched (n = 4896) 1-1 orthologous protein-coding genes. **f,** Screenshot of DevSeqPlant, a web-based interactive analysis and visualisation platform for plant evolutionary transcriptomics (https://www.devseqplant.org).

**Extended Data Table 1.** Plant samples for profiling. Sample size denotes number of collected samples/number of plants.

| ID | Species | Tissue | Description | Age | Substrate | Sample Size |
| --- | --- | --- | --- | --- | --- | --- |
| 10-12 | *A.thaliana* | root | root tip (top 0.5-1mm) | 5d | ½ MS agar | 90 |
| 13-15 | *A.thaliana* | root | maturation zone (2-3mm) | 5d | ½ MS agar | 90 |
| 106-108 | *A.thaliana* | root | root, whole root (3mm) | 5d | ½ MS agar | 45 |
| 1-3 | *A.thaliana* | root | root, whole root | 7d | soil | 50 |
| 7-9 | *A.thaliana* | root | root, whole root | 14d | soil | 20 |
| 4-6 | *A.thaliana* | root | root, whole root | 21d | soil | 30 |
| 16-18 | *A.thaliana* | hypocotyl | hypocotyl | 10d | soil | 35 |
| 22-24 | *A.thaliana* | stem | 3rd internode (0.5cm) | 24d | soil | 25 |
| 19-21 | *A.thaliana* | stem | 2nd internode (1cm piece) | 24d | soil | 25 |
| 25-27 | *A.thaliana* | stem | 1st internode (1cm piece) | 28d+ | soil | 20 |
| 28-30 | *A.thaliana* | leaf | cotyledons | 7d | soil | 50/25 |
| 31-33 | *A.thaliana* | leaf | leaves 1+2 | 7d | soil | 60/30 |
| 34-36 | *A.thaliana* | leaf | leaves 1+2 | 10d | soil | 40/20 |
| 40-42 | *A.thaliana* | leaf | leaves 1+2 - petiole | 10d | soil | 60/30 |
| 43-45 | *A.thaliana* | leaf | leaves 1+2 - leaf tip | 10d | soil | 60/30 |
| 37-39 | *A.thaliana* | leaf | leaves 5+6 | 17d | soil | 40/20 |
| 286-288 | *A.thaliana* | leaf | leaves 9+10 | 27d | soil | 40/20 |
| 46-48 | *A.thaliana* | leaf | leaves senescing | 35d | soil | 30 |
| 118-120 | *A.thaliana* | leaf | cauline leaves | 24d | soil | 30 |
| 115-117 | *A.thaliana* | apex | apex, vegetative | 7d | soil | 40 |
| 49-51 | *A.thaliana* | apex | apex, vegetative | 10d | soil | 40 |
| 52-54 | *A.thaliana* | apex | apex, veg, before bolting | 14d | soil | 35 |
| 55-57 | *A.thaliana* | apex | apex inflorescence | 21d | soil | 30 |
| 58-60 | *A.thaliana* | apex | apex inflorescence, *clv1-8* | 21d+ | soil | 30 |
| 61-63 | *A.thaliana* | apex | apex inflorescence | 28d | soil | 30 |
| 64-66 | *A.thaliana* | flowers | flower stage 9 | 21d+ | soil | 100/25 |
| 67-69 | *A.thaliana* | flowers | flower stage 10/11 | 21d+ | soil | 100/25 |
| 70-72 | *A.thaliana* | flowers | flower stage 12 | 21d+ | soil | 75/25 |
| 73-75 | *A.thaliana* | flowers | flower stage 15 | 21d+ | soil | 60/30 |

| ID | Species | Tissue | Description | Age | Subst rate | Sample Size |
| --- | --- | --- | --- | --- | --- | --- |
| 76-78 | *A.thaliana* | floral organs | sepals stage 12 | 21d+ | soil | 140/35 |
| 88-90 | *A.thaliana* | floral organs | sepals stage 15 | 21d+ | soil | 100/25 |
| 79-81 | *A.thaliana* | floral organs | petals stage 12 | 21d+ | soil | 140/35 |
| 91-93 | *A.thaliana* | floral organs | petals stage 15 | 21d+ | soil | 100/25 |
| 82-84 | *A.thaliana* | floral organs | stamen stage 12 | 21d+ | soil | 200/35 |
| 94-96 | *A.thaliana* | floral organs | stamen stage 15 | 21d+ | soil | 150/25 |
| 109-111 | *A.thaliana* | floral organs | mature pollen | 21d+ | soil | 50 plants |
| 283-285 | *A.thaliana* | floral organs | carpel stage 12 (early) | 21d+ | soil | 80/30 |
| 85-87 | *A.thaliana* | floral organs | carpel stage 12 (late) | 21d+ | soil | 50/25 |
| 97-99 | *A.thaliana* | floral organs | carpel stage 15 (w/ seeds) | 21d+ | soil | 45/45 |
| 100-102 | *A.thaliana* | fruit/seeds | fruit stage 16 w/seeds | 28d+ | soil | 30 |
| 103-105 | *A.thaliana* | fruit/seeds | fruit stage 17a w/seeds | 28d+ | soil | 30 |
| 289-291 | *A.thaliana* | fruit/seeds | seeds - fruit stage 16 | 28d+ | soil | 40 siliques |
| 292-294 | *A.thaliana* | fruit/seeds | seeds - fruit stage 17a | 28d+ | soil | 40 siliques |
| 112-114 | *A.thaliana* | fruit/seeds | seeds - fruit stage 18 | 28d+ | soil | 40 siliques |
| 121-123 | *A.lyrata* | root | root, whole root (3mm) | 6d | ½ MS | 70 |
| 124-126 | *A.lyrata* | hypocotyl | hypocotyl | 12d | soil | 30 |
| 127-129 | *A.lyrata* | leaf | leaves 1+2 | 10d | soil | 60/30 |
| 130-132 | *A.lyrata* | apex | apex, vegetative | 11d | soil | 35 |
| 133-135 | *A.lyrata* | apex | apex, inflorescence | 8w/10w/13d | soil | 35 |
| 136-138 | *A.lyrata* | flower | flower stg. 12 (mid-stage) | 8w/10w/15d | soil | 60/20 |
| 139-141 | *A.lyrata* | floral organ | mature pollen | 8w/10w/25d | soil | 30 plants |
| 247-249 | *A.lyrata* | floral organ | carpel stg. 12 (early/mid) | 8w/10w/17d | soil | 60/20 |
| 295-297 | *A.lyrata* | floral organ | stamen stg. 11 | 8w/10w/25d | soil | 150/30 |
| 298-300 | *A.lyrata* | floral organ | stamen stg 12 (early stage) | 8w/10w/23d | soil | 150/20 |
| 304-306 | *A.lyrata* | floral organ | stamen stg- 12 (mid-stage) | 8w/10w/24d | soil | 220/35 |
| 301-303 | *A.lyrata* | floral organ | stamen stg- 12 (late stage) | 8w/10w/21d | soil | 250/25 |
| 142-144 | *C.rubella* | root | root, whole root (3mm) | 4d | ½ MS | 70 |
| 145-147 | *C.rubella* | hypocotyl | hypocotyl | 9d | soil | 35 |

**Extended Data Table 1. Plant samples for profiling.**
| ID | Species | Tissue | Description | Age | Substrate | Sample Size |
| --- | --- | --- | --- | --- | --- | --- |
| 148-150 | *C.rubella* | leaf | leaves 1+2 | 7d | soil | 60/30 |
| 151-153 | *C.rubella* | apex | apex, vegetative | 7d | soil | 35 |
| 154-156 | *C.rubella* | apex | apex, inflorescence | 6w/7w/17d | soil | 30 |
| 157-159 | *C.rubella* | flower | flower stage 12 (mid) | 6w/7w/20d | soil | 75/25 |
| 160-162 | *C.rubella* | floral organ | mature pollen | 6w/7w/22d | soil | 50 plants |
| 250-252 | *C.rubella* | floral organ | carpel stage 12 | 6w/7w/21d | soil | 65/25 |
| 268-270 | *C.rubella* | floral organ | stamen stage 12 | 6w/7w/20d | soil | 220/25 |
| 163-165 | *E.salsugineum* | root | whole root (3mm) | 6d | ½ MS | 100 |
| 166-168 | *E.salsugineum* | hypocotyl | hypocotyl | 12d | soil | 35 |
| 169-171 | *E.salsugineum* | leaf | leaves 1+2 | 9d | soil | 80/40 |
| 172-174 | *E.salsugineum* | apex | apex, vegetative | 9d | soil | 40 |
| 175-177 | *E.salsugineum* | apex | apex, inflorescence | 7w/8w/12d | soil | 30 |
| 178-180 | *E.salsugineum* | flower | flower stage 12 | 7w/8w/15d | soil | 75/25 |
| 181-183 | *E.salsugineum* | floral organ | mature pollen | 7w/8w/17d | soil | 50 plants |
| 253-255 | *E.salsugineum* | floral organ | carpel stage 12 | 7w/8w/18d | soil | 75/25 |
| 271-273 | *E.salsugineum* | floral organ | stamen stage 12 | 7w/8w/17d | soil | 180/30 |
| 184-186 | *T.hassleriana* | root | whole root (3mm) | 5d | ½ MS | 30 |
| 187-189 | *T.hassleriana* | hypocotyl | hypocotyl | 11d | soil | 25 |
| 190-192 | *T.hassleriana* | leaf | leaves 1+2 | 11d | soil | 50/25 |
| 193-195 | *T.hassleriana* | apex | apex, vegetative | 11d | soil | 30 |
| 196-198 | *T.hassleriana* | apex | apex, inflorescence | 11w | soil | 20 |
| 199-201 | *T.hassleriana* | flower | flower stg.12 equiv. | 11w | soil | 75/25 |
| 202-204 | *T.hassleriana* | floral organ | mature pollen | 11w | soil | 50 plants |
| 256-258 | *T.hassleriana* | floral organ | carpel stage 12 | 11w | soil | 75/25 |
| 274-276 | *T.hassleriana* | floral organ | stamen stage 12 | 11w | soil | 150/25 |
| 205-207 | *M.truncatula* | root | whole root (3mm) | 4d | ½ MS | 40 |
| 208-210 | *M.truncatula* | hypocotyl | hypocotyl | 8d | soil | 30 |
| 211-213 | *M.truncatula* | leaf | leaf 2 | 7d | soil | 25 |
| 214-216 | *M.truncatula* | apex | apex, vegetative | 6d | soil | 30 |
| 217-219 | *M.truncatula* | apex | inflorescence meristem including I1, I2 and F, Br, L, Stp primordia | 7w | soil | 30 |
| 220-222 | *M.truncatula* | flower | flower stage 8 | 7w | soil | 45/25 |
| 223-225 | *M.truncatula* | floral organ | mature pollen | 7w | soil | 20 plants |
| 259-261 | *M.truncatula* | floral organ | carpel flower stage 8 | 7w | soil | 50/25 |
| 277-279 | *M.truncatula* | floral organ | stamen flower stage 8 | 7w | soil | 200/20 |
| 226-228 | *B.distachyon* | root | whole root (3mm) | 4d | ½ MS | 25 |
| 229-231 | *B.distachyon* | mesocotyl | mesocotyl | 7d | soil | 35 |
| 232-234 | *B.distachyon* | leaf | leaf 1 | 4d | soil | 30 |
| 235-237 | *B.distachyon* | apex | apex, vegetative | 5d | soil | 35 |
| 238-240 | *B.distachyon* | apex | Lateral spikelet meristem (“naked-stage” to “awn-initiation” stage) | 18-20d | soil | 35 |
| 241-243 | *B.distachyon* | flower | floret with anthers immediately before dehiscence from basal lateral spikelet | 32d | soil | 60 |
| 244-246 | *B.distachyon* | floral organ | mature pollen from basal lateral spikelet | 32d | soil | 40 plants |
| 262-264 | *B.distachyon* | floral organ | carpels “stage 12 equivalent” from basal lateral spikelet | 32d | soil | 50/25 |
| 280-282 | *B.distachyon* | floral organ | stamen “stage 12 equivalent” from basal lateral spikelet | 32d | soil | 100/25 |

